# The METTL5-TRMT112 *N^6^*-methyladenosine methyltransferase complex regulates metabolism and development via translation

**DOI:** 10.1101/2021.05.18.444567

**Authors:** Caraline Sepich-Poore, Zhong Zheng, Emily Schmitt, Kailong Wen, Zijie Scott Zhang, Xiao-Long Cui, Qing Dai, Allen C. Zhu, Linda Zhang, Arantxa Sanchez Castillo, Xiaoxi Zhuang, Chuan He, Sigrid Nachtergaele

## Abstract

Ribosomal RNAs (rRNAs) have long been known to carry modifications, including numerous sites of 2’*O*-methylation and pseudouridylation, as well as *N*^6^-methyladenosine (m^6^A), and *N*^6,6-^dimethyladenosine. While the functions of many of these modifications are unclear, some are highly conserved and occur in regions of the ribosome critical for mRNA decoding. Both 28S rRNA and 18S rRNA carry m^6^A, and while ZCCHC4 has been identified as the methyltransferase responsible for the 28S rRNA m^6^A site, the methyltransferase responsible for the 18S rRNA m^6^A site has remained uncharacterized until recently. Here, we show that the METTL5-TRMT112 complex is the methyltransferase responsible for installing m^6^A at position 1832 of human 18S rRNA. TRMT112 is required for the metabolic stability of METTL5, and human METTL5 mutations associated with microcephaly and intellectual disability disrupt this interaction. Loss of METTL5 in human cancer lines alters the translation of transcripts associated with mitochondrial biogenesis and function. *Mettl5* knockout mice display reduced body size and evidence of metabolic defects. This m^6^A site is located on the 3’ end of 18S rRNA, which may become surface-exposed under some circumstances and thus may play a regulatory role in translation of specific transcripts. While recent work has focused heavily on m^6^A modifications in mRNA and its roles in mRNA processing and translation, deorphanizing putative methyltransferase enzymes is revealing previously unappreciated regulatory roles for m^6^A in noncoding RNAs.

## INTRODUCTION

Chemical modifications on RNA are a critical facet of gene expression regulation. Historically, modifications on tRNA and rRNA have been thought to have high-stoichiometry and be relatively static, while work in the last decade on mRNA modifications suggests they are often sub-stoichiometric and more dynamic (1). rRNA is heavily modified with numerous chemical marks, including pseudouridine, 2’*O*-methylation (2’*O*Me), *N*^7^-methylguanosine (m^7^G), *N*^1^-methyladenosine (m^1^A), *N*^6^-methyladenosine (m^6^A), and *N*^6,6^-methyladenosine (m^6,6^A) (2-4). While these modifications are thought to play critical structural roles and many of the regulatory enzymes have been identified, it is often difficult to assign specific functions to individual modifications due to the numerous interactions between the three rRNAs and 80+ protein components that form the ribosome. While some modifications play structural roles in ribosome assembly, others may regulate translation of specific transcripts. Disruption of rRNA modification processes has been implicated in a class of developmental disorders called ribosomopathies (5-8). Interestingly, though the ribosome is ubiquitously essential for translating protein, ribosomopathies often manifest as tissue-specific disorders, the molecular mechanisms of which we do not understand in many cases (9).

The METTL protein family is a class of S-adenosyl-methionine-dependent methyltransferases, with over thirty family members that methylate DNA (10), RNA (11-14), and protein (15,16) substrates. Some, such as METTL3, METTL14, and METTL16, have well-characterized functions as RNA m^6^A methyltransferases (11,12), but others remain poorly understood. Notably, mutations in many of these enzymes, including those whose functions are poorly understood, have been implicated in human diseases such as developmental abnormalities and cancers (17-19). Revealing METTL protein substrate specificity, activity, and function are critical first steps towards understanding how mutations in these enzymes cause human disease. More specifically, mutations in METTL5 have been implicated in developmental abnormalities including microcephaly, intellectual disabilities, and attention deficit hyperactivity disorder (ADHD), but until recently very little was known about METTL5 function (18,20). We were intrigued by this connection when we came across METTL5 via proteomics experiments aimed at identifying novel proteins with methyltransferase activity.

While this work was in progress, multiple studies revealed that a complex containing METTL5 and TRMT112 m^6^A-methylates 18S rRNA in flies, mice, and humans (21-24). Mettl5 in *C. elegans* also carries out this function, but not in the context of a complex with TRMT112, as *C. elegans* lack a TRMT112 homologue (25,26). Interestingly, the role of METTL5 in protein translation differs across the different model organisms studied. Consistent with these reports, we find that METTL5 forms a complex with TRMT112 to m^6^A methylate 18S rRNA in human cell lines. We further find that TRMT112 is critical for stabilizing METTL5 at the protein level, and that depletion of TRMT112 is sufficient to reduce METTL5 protein levels. Catalytically inactive METTL5 mutants can retain this critical association with TRMT112, but METTL5 mutations derived from human patients dramatically reduce this interaction. While we did not see global changes in protein translation upon METTL5 depletion, we did find evidence of dysregulated translation of specific genes. To complement our cellular studies, we generated *Mettl5* knockout mice, and validated the loss of 18S rRNA m^6^A1832 in tissues from these mice. Consistent with Ignatova *et al*. and Wang *et al*., we observe smaller body size in our *Mettl5* KO mouse model (22,27). We do not observe significant behavioral changes that have been previously described, but through visual observation and RNA sequencing of mouse tissues, we do find evidence of metabolic defects that have not yet been reported. We propose that METTL5 may regulate the translation of transcripts associated with lipid metabolism, resulting in metabolic dysregulation that may play a role in the developmental phenotypes seen in human patients.

## RESULTS

### METTL5 is an m^6^A methyltransferase stabilized by TRMT112

METTL5 is a member of the METTL family of S-adenosyl-methionine-dependent methyltransferases, which is not found in yeast but conserved in higher eukaryotes ranging from *C. elegans*, to mice, to humans. While the activities and functions of some METTL proteins have been elucidated over the last decade (11-14), many remain poorly understood. RNA methyltransferases, particularly the METTL3-METTL14 complex, have been demonstrated to have numerous cellular functions through their methyltransferase activity, which raises the question as to how less well-characterized METTL proteins might regulate cellular processes. METTL5 drew our attention in the course of a biochemical screen which was aimed at identifying novel RNA methyltransferase activity through biochemical fractionation followed by a liquid chromatography coupled to tandem mass spectrometry (LC-MS/MS)-based assay (data not shown). We surveyed commonly used cancer cell culture lines by western blot and found that HeLa, HEL, and K562 cells showed relatively higher METTL5 expression (Figure 1A). HeLa cells are the only adherent line of these three and are easily transfected, making biochemical and imaging studies straightforward. Thus, we generated METTL5 knockout (KO) HeLa lines using CRISPR-Cas9 and two sets of two guide RNAs to generate deletions across two different exons (Supp. Fig. 1A). After puromycin selection, single clones were isolated, expanded, and tested for METTL5 expression by western blot (Figure 1B; Supp. Fig. 1A). Through this process, we isolated both KO lines and clones in which METTL5 expression was unaffected (e.g., clones 4 and 6), which serve as controls throughout this work.

**Figure 1.**
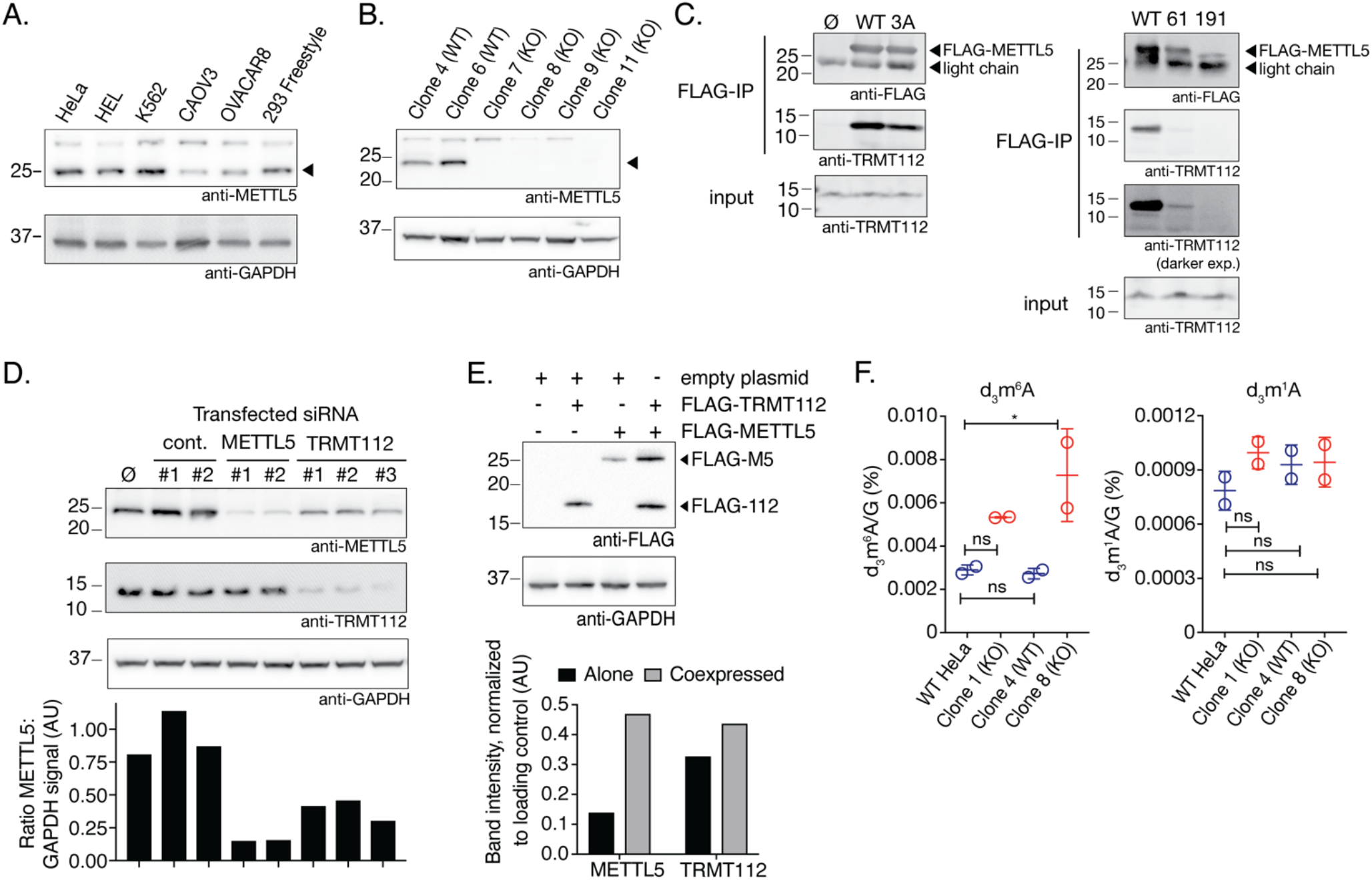
METTL5 is an m^6^A methyltransferase stabilized by TRMT112. **(A)** METTL5 expression in cell lines as evaluated by western blot (top) with anti-GAPDH loading control (bottom). Arrowhead signifies METTL5 band. **(B)** Western blot analysis of METTL5 levels in wild type (WT) and knockout (KO) HeLa samples, each expanded from a single isolated clone (top) with anti-GAPDH loading control (bottom). Arrowhead signifies METTL5 band. **(C)** Co-immunoprecipitations of TRMT112 with FLAG-METTL5. Left: Anti-FLAG (top) and anti-TRMT112 (middle) western blots in anti-FLAG immunoprecipitation samples. Input loading control shown on bottom. Ø, untransfected HeLa cells; WT, HeLa transfected with FLAG-METTL5; 3A, HeLa transfected with FLAG-METTL5-3A. Right: Anti-FLAG (top) and anti-TRMT112 (middle) western blots in anti-FLAG immunoprecipitation samples. Input loading control shown on bottom. WT, HeLa transfected FLAG-METTL5-WT; 61, HeLa transfected with FLAG-METT5^G61D^; 191, HeLa transfected with FLAG-METTL5^K191Vfs*10^. **(D)** Top: western blots for METTL5, TRMT112, and GAPDH (loading control) levels upon siRNA knockdown for METTL5 and TRMT112. Ø, untransfected HeLa cells; cont. HeLa transfected with two different non-targeting control siRNAs; METTL5, HeLa transfected with two different siRNAs targeting METTL5; TRMT112, HeLa transfected with three different siRNAs targeting TRMT112. Bottom: ratio of METTL5 to GAPDH signal intensity as quantified by Fiji software. **(E)** Top: western blot analysis of anti-FLAG immunoprecipitations from HeLa transfected with labeled combinations of vectors, with GAPDH loading control. Bottom: Band intensity normalized to GAPDH band intensity, as quantified by Fiji software. **(F)** LC-MS/MS analysis of d_3_m^6^A (left) and d_3_m^1^A (right) levels normalized to guanosine from *in vitro* methyltransferase reactions performed on total RNA isolated from METTL5-WT and METTL5-KO HeLa cells. n=2 replicate reactions, mean and s.e.m. plotted, analyzed by one-way ANOVA, comparing all samples to HeLa WT with Dunnett’s test for multiple comparisons. ns: not significant, * p <0.05

Based on the presence of an NPPF motif, similar to other motifs found in m^6^A RNA methyltransferases (e.g., DPPW in METTL3), we hypothesized that METTL5 may be an m^6^A methyltransferase. However, through our efforts to use *in vitro* methylation assays with tagged, overexpressed METTL5 protein to validate its activity, we noted lower-than-typical yields of purified METTL5 protein from both bacterial and mammalian expression systems, and low modification fractions in our *in vitro* assays (data not shown). Since methyltransferases such as METTL3-METTL14 function as a complex, this prompted us to perform proteomic analysis of METTL5 binding proteins using FLAG-METTL5 expressed in FreeStyle 293-F cells. TRMT112 was a particularly intriguing candidate binding protein (Supp. Fig. 1B), because it is known to bind and regulate other methyltransferases such as AlkBH8 (a tRNA methyltransferase) (28) and HemK2 (a protein methyltransferase) (29). Indeed, we found a direct interaction between METTL5 and TRMT112 (Figure 1C) (21). Mutation of the METTL5 NPP to AAA (amino acids 127-129, abbreviated METTL5-3A), predicted to abrogate its m^6^A methyltransferase activity, slightly weakened but did not completely disrupt this interaction. Mutations in METTL5 have also been found in human patients with intellectual disability and microcephaly (18,20). We introduced two of these human variants, G61D and K191Vfs*10, into our FLAG-METTL5 construct. While FLAG-METTL5-G61D expressed at similar levels to METTL5-WT, the interaction with TRMT112 was significantly compromised, potentially explained by structural changes introduced by disrupting flexibility of a loop region (Figure 1C, Supp. Fig. 1C). FLAG-METTL5-K191Vfs*10 expressed at much lower levels, suggesting that this truncated protein may not fold properly (Figure 1C). Based on a previous report, the third variant, METTL5-R115Nfs*19, expresses only at very low levels, so we did not test its expression or interaction with TRMT112 (20).

The interaction with TRMT112 stabilizes METTL5 protein, as siRNA knockdown of TRMT112 reduced expression of METTL5 to approximately half, relative to negative controls (Figure 1D), though not to the same extent as siRNA knockdown of METTL5 directly. The stabilization effect occurs at the protein level, as METTL5 transcript levels are unaffected by TRMT112 knockdown (Supp. Fig. 1D). Conversely, coexpression of FLAG-METTL5 with FLAG-TRMT112 in HeLa cells significantly increased FLAG-METTL5 protein expression relative to solely expressing FLAG-METTL5 (Figure 1E). Coexpression of FLAG-TRMT112 with FLAG-METTL5 substantially increased protein expression yields from FreeStyle 293-F cells, allowing us to purify FLAG-METTL5-FLAG-TRMT112 complex for *in vitro* methyltransferase assays. Using deuterated S-adenosyl methionine (d_3_SAM), we performed *in vitro* methyltransferase assays with co-purified FLAG-METTL5-FLAG-TRMT112, using total RNA isolated from METTL5-WT and METTL5-KO HeLa cells. Analysis of deuterated, methylated nucleosides by LC-MS/MS revealed the appearance of d_3_m^6^A in all four samples (Figure 1F). RNA from METTL5-KO cells accumulated approximately double the amount of d_3_m^6^A as RNA isolated from METTL5-WT cells, consistent with the idea that total RNA from METTL5-KO cells has less m^6^A to begin with. Parallel measurements in these same samples showed 10-fold lower levels of d_3_m^1^A that remained consistent across all samples. Taken together, these results suggest that METTL5 is an m^6^A RNA methyltransferase that requires TRMT112 for stability, as also corroborated by van Tran *et al*. (21).

### 18S rRNA is a major substrate of the METTL5-TRMT112 complex

To begin to understand the function of the METTL5-TRMT112 complex in cells, we first sought to identify its subcellular distribution. Fluorescence microscopy using antibodies targeting endogenous METTL5 revealed nuclear puncta that colocalized with the nucleolar protein fibrillarin, but also had a more diffuse staining pattern in the cytoplasm (Figure 2A). This distribution was also verified using biochemical fractionation, which revealed that, while there was no detectable METTL5 associated with chromatin, there is a small pool of nuclear METTL5, with the majority of the protein localized to the cytoplasm (Figure 2B). This subcellular localization is particularly intriguing given the large difference observed in d_3_m^6^A methylation of total RNA isolated from WT and KO cells (Figure 1F), suggesting a relatively large pool of RNA to be methylated by METTL5-TRMT112. The nucleolus is a critical hub for the processing and assembly of ribosome components, and rRNAs are known to be m^6^A methylated. Both 28S and 18S rRNAs have an m^6^A site (3), and while ZCCHC4 has been identified as the methyltransferase for 28S rRNA A4220 (30), the methyltransferase for 18S rRNA was unknown when we initiated these studies, and was first reported in van Tran *et al*. (21).

**Figure 2.**
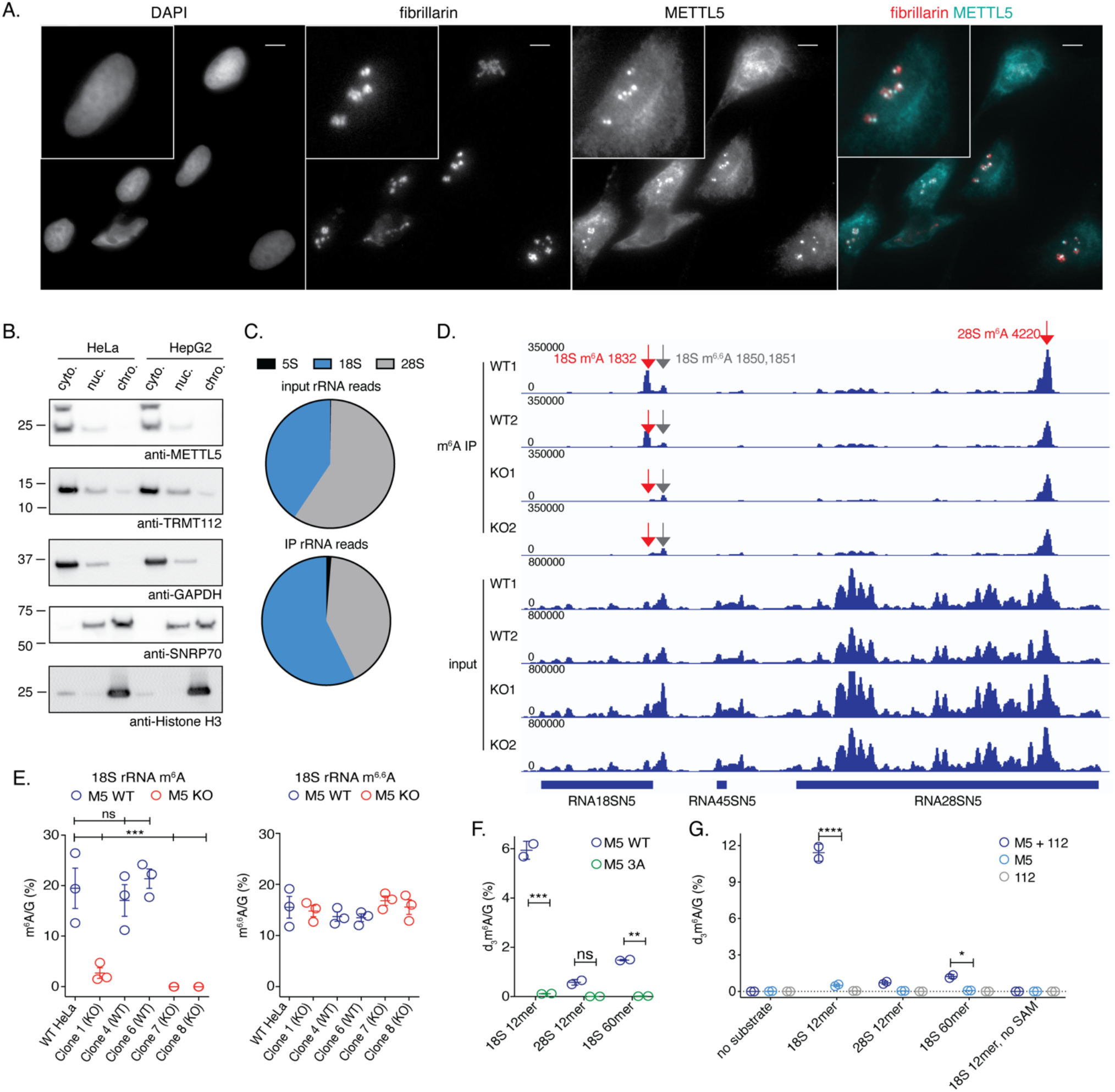
METTL5 is predominantly cytoplasmic and methylates 18S rRNA. **(A)** Fluorescence microscopy images of HeLa cells stained with DAPI (nuclear stain), anti-fibrillarin antibody (nucleolar marker), and anti-METTL5 antibody. Right panel shows merged image. Scale bar: 10μm. **(B)** Localization of METTL5 and TRMT112 in chromatin-associated, nuclear, and cytoplasmic cellular fractions in HeLa and HepG2 by western blot. Loading and fractionation controls: GAPDH (cytoplasmic), SNRP70 (nuclear), Histone H3 (chromatin-associated). **(C)** Pie charts of reads aligned to 5S, 18S, or 28S rDNA from HeLa input and METTL5 CLIP samples. **(D)** Visualization of reads aligned to the pre-45S locus from input and anti-m^6^A immunoprecipitated Me-RIP-seq samples from wild type and METTL5 knockout HeLa cells, adapted from IGV. **(E)** Levels of m^6^A (left) and m^6,6^A (right), normalized to G, obtained by LC-MS/MS of 40-nt probe-purified segments of 18S rRNA surrounding the m^6^A 1832 site from HeLa METTL5-WT and METTL5-KO cells. n=3 reactions per condition, mean and s.e.m. plotted. Comparisons of all samples to the WT control were tested by one-way ANOVA with Dunnett’s test for multiple comparisons. **(F)** LC-MS/MS analysis of d_3_m^6^A levels normalized to G from *in vitro* methyltransferase reactions performed on 18S and 28S-mimicking probes with TRMT112 and either wild type (WT) or catalytic mutant (3A) METTL5. **(G)** LC-MS/MS analysis of d_3_m^6^A levels normalized to G from *in vitro* methyltransferase reactions performed on 18S and 28S probes with METTL5 alone, TRMT112 alone, or METTL5-TRMT112 complex. (F) and (G) were analyzed using two-way ANOVA to compare WT with 3A (with Sidak test for multiple comparisons, F) and METTL5 with METTL5-TRMT112 (with Tukey test for multiple comparisons, G). n=2 reactions per condition, mean and s.e.m. * p<0.05, ** p<0.01, *** p<0.005. **** p<0.0001 (all other comparisons not statistically significant [ns], not indicated).

To identify RNA substrates of METTL5, we performed crosslinking-assisted immunoprecipitation (CLIP) of FLAG-tagged METTL5 followed by high throughput sequencing of bound RNAs. To maximize the chances of success, we also incorporated 4-thiouridine to facilitate crosslinking (31) (see ‘Materials and Methods’ for a detailed protocol). High-throughput sequencing revealed a slight enrichment for 18S rRNA transcripts in the immunoprecipitate (IP) relative to input (Figure 2C), as well as other enriched transcripts (Supp. Fig. 2A). m^6^A-seq in METTL5-WT and -KO HeLa cells showed loss of an m^6^A peak near adenosine 1832 in 18S rRNA while the adjacent m^6,6^A1850 and m^6,6^A1851 sites remained unchanged (Figure 2D), supporting the hypothesis that 18S rRNA is a major substrate of the METTL5-TRMT112 complex. We applied biotinylated DNA probes complementary to the region of 18S rRNA containing both m^6^A1832 and the neighboring m^6,6^A1850,1851 sites to capture and purify these fragments from METTL5-WT and -KO cells for LC-MS/MS analysis. Results confirmed that METTL5-TRMT112 methylates 18S rRNA A1832, as this fragment showed dramatically lower levels of m^6^A in all KO lines tested, relative to WT (Figure 2E, left panel). In contrast, the nearby m^6,6^A sites showed almost no change among these cell lines (Figure 2E, right panel), demonstrating that 18S rRNA processing is not dramatically altered, and suggesting that these modifications are regulated independently.

To verify that METTL5-TRMT112 is not only necessary, but also sufficient for deposition of 18S rRNA m^6^A1832, we expressed and purified FLAG-METTL5-WT and catalytically inactive FLAG-METTL5-3A with FLAG-TRMT112 in FreeStyle 293-F cells and tested their activity on RNA probes containing 18S or 28S rRNA sequences, using LC-MS/MS to measure d_3_m^6^A levels. Neither FLAG-METTL5-WT nor FLAG-METTL5-3A complexes could efficiently methylate a 28S rRNA 12-mer (Figure 2F), suggesting specificity for 18S rRNA sequence. Only METTL5-WT-TRMT112 complexes could effectively methylate 18S rRNA 12-mer or 60-mer probes, while the 3A mutant showed nearly undetectable activity. With this highly effective 18S rRNA 12mer substrate in hand, we performed a more detailed assessment of the individual contributions of METTL5 and TRMT112 in methyltransferase activity. While copurified FLAG-METTL5 and FLAG-TRMT112 could effectively methylate 18S rRNA probes, this activity was undetectable when either protein was purified individually, or when d_3_SAM was left out of the reaction (Figure 2G). The loss of METTL5 activity in the absence of TRMT112 is likely the result of poorly folded METTL5 protein, as suggested by increased stability of METTL5 in the presence of TRMT112 (Figures 1D, 1E). Taken together, our results are consistent with recent reports (21,22,24) that the METTL5-TRMT112 complex m^6^A methylates 18S rRNA and that the NPPF motif is critical for its catalytic activity.

To assess whether METTL5-TRMT112 may have other RNA substrates, we delved more deeply into the other transcripts enriched in our METTL5 CLIP experiment (Supp. Fig. 2A,B), which included both coding and non-coding RNAs. Cross-referencing these METTL5-bound RNAs with differentially methylated transcripts from our m^6^A-seq experiment (Supp. Fig. 2C) revealed only 25 overlapping transcripts (Supp. Fig. 2D). While the most prominent motif in the m^6^A peaks overall was GGACU, suggesting that most m^6^A peaks were METTL3/METTL14 dependent (11), the 25 transcripts in common between the two datasets showed enrichment for UAA, the motif containing the m^6^A site in 18S rRNA. Thus, though it has been reported that 18S rRNA is the only METTL5-TRMT112 substrate (21), it remains possible that other targets may exist. Notably, the coding transcripts identified in our CLIP experiment are enriched for genes involved in mitochondrial biogenesis and function (Supp. Fig. 2B). RNA-seq analysis of differentially expressed transcripts in METTL5-WT and -KO HeLa cells also revealed enrichment for small molecule transport and lipid and cholesterol biosynthesis pathways (Supp. Fig. 2E). These preliminary connections to metabolism and lipid biosynthesis, both liver-based functions, led us to generate HepG2 METTL5-KO cell lines, which may reflect gene expression pathways in the liver (Supp. Fig. 2F, 2G) more closely.

### METTL5 regulates translation of a subset of transcripts

Through the course of our experiments with both HeLa and HepG2 cells, we noted that METTL5-KO cells tended to grow more slowly than the corresponding METTL5-WT cells. To measure this difference directly, cell growth curves were monitored over the course of 96 hours using DNA-dye-based CyQuant cell proliferation assays. In both HeLa and HepG2 cells, METTL5 KO cell lines tended to grow more slowly than wild type cells, with a small but consistent difference in growth rate (Figure 3A, Supp. Fig. 3A). rRNAs are heavily modified with numerous chemical groups which regulate ribosome biogenesis and function, and play diverse roles in gene expression regulation. The slight growth defect we observed suggested that METTL5 is likely not essential for ribosome biogenesis (21), but that it may regulate the translation of a subset of transcripts that collectively slows cell proliferation. Consistent with this, polysome profiling in both HeLa (Figure 3B, 3C) and HepG2 cells (Supp. Fig. 3B) shows no notable changes in global translation. We note that the effect of METTL5 depletion on global translation has differed across recently published reports on METTL5 function, especially across different cell lines (21-25).

**Figure 3.**
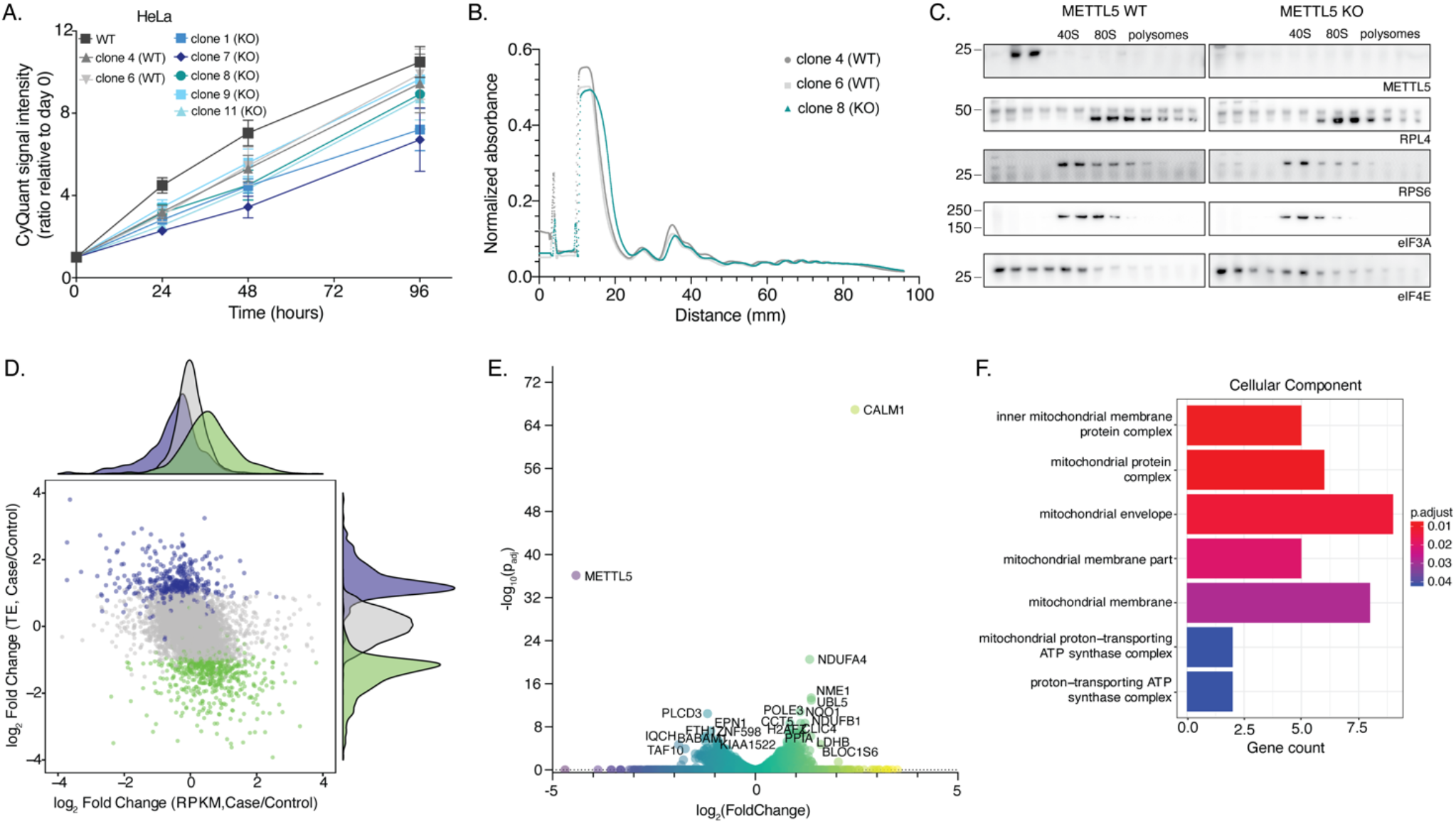
METTL5 affects cell proliferation and metabolism through regulation of translation of specific transcripts. **(A)** Cell proliferation of HeLa METTL5-WT and METTL5-KO cells over time as measured by CyQuant assay. Means (squares) and SDs (bars) are indicated for 4 replicate wells per condition. **(B)** Polysome profiles from HeLa METTL5-WT and METTL5-KO cells as measured by normalized absorbance over a 5-50% sucrose gradient. **(C)** Western blots of polysome fractions from panel B for METTL5, RPL4 (large ribosomal subunit marker), RPS6 (small ribosomal subunit marker), eIF3A, and eIF4E (translation initiation factors). **(D)** Effects of METTL5 knockout on translation and transcription as displayed by log_2_(fold change) of translation efficiency (TE) versus log_2_(fold change) of reads per kilobase of transcript per million mapped reads (RPKM) from ribosome profiling input and ribosome-protected fragment sequencing of HepG2 cells. Case = METTL5 knockout, Control = wild type. **(E)** Volcano plot of the negative log_10_ of adjusted P-value versus log_2_(fold change) of reads from ribosome profiling of HepG2 METTL5-KO versus METTL5-WT cells. **(F)** Cellular component gene ontology (GO) functional enrichment analysis for translationally upregulated transcripts in HepG2 METTL5-KO compared to METTL5-WT cells. D and F are modified from output of RiboToolKit (40).

To identify specific transcripts whose regulation may be disrupted by loss of METTL5, we sequenced ribosome protected fragments. At the global level, ribosome profiling experiments in HepG2 cells revealed more significant changes at the level of transcript translation than transcript abundance. In comparing METTL5-WT versus -KO cells, translation efficiency (TE) showed greater variance than transcript abundance (measured as reads per kilobase of transcript per million mapped reads, RPKM), suggesting that gene expression changes in METTL5 KO cells may occur largely at the level of transcript translation rather than at the transcriptional level (Figure 3D). Interestingly, transcripts upregulated in METTL5-KO cells were enriched for transcriptional regulators and cholesterol and lipid biosynthetic processes (Supp. Fig. 2E).

Sequencing of ribosome-protected fragments revealed several differentially translated transcripts in METTL5 KO cells (Figure 3E), the two most striking being METTL5, which was expectedly significantly downregulated in METTL5-KO cells, and CALM1, a calcium binding protein known to regulate cell proliferation and growth. Upon inspecting ribosome-protected fragment and input reads aligned to CALM1 in IGV, we noted similar transcript levels in both WT and KO cells, but greatly increased ribosome occupancy in KO cells, especially in exon 4 (Supp. Fig. 3C). In addition, numerous genes involved in the biogenesis and regulation of mitochondria were also significantly upregulated in METTL5 KO cells (Figure 3F). Overall, although our ribosome profiling data showed that a subset of transcripts were affected more than others, it did not point to a clear mechanism explaining the preference for certain transcripts. Annotation of translated ORFs revealed higher levels of internal and novel ORFs, suggesting that dysregulated translation may be the result of frameshifting or translation of ordinarily non-coding transcripts (Supp. Fig. 3D). While there was slight dysregulation across codons due to METTL5 depletion, no single codon or amino acid stood out as being particularly affected in terms of occupancy at the P-site, which is near 18S m^6^A1832 (Supp. Fig. 3E).

### *Mettl5* KO mice demonstrate growth and metabolic changes

To assess METTL5 function at the level of a whole organism, we generated *Mettl5* knockout mice (*Mettl5*^-/-^) by disrupting exon 2 of *Mettl5* with CRISPR-Cas9 (Supp. Fig. 4A). *Mettl5* expression was undetectable in these mice by qPCR and greatly diminished by RNA-seq (Supp. Fig. 4B,C). Critically, m^6^A levels on 18S rRNA isolated directly from the brains and livers of these mice were abolished, while the neighboring m^6,6^A site remained stable, consistent with our findings in METTL5-KO HeLa and HepG2 cells (Figure 4A). Consistent with Ignatova *et al*. (22), we observe that *Mettl5* KO mice are subviable and saw fewer than expected *Mettl5*^-/-^ mice from both HET/HET (Figure 4B) and HET/KO (Supp. Fig. 4D) breeding pairs. It was also immediately, visibly apparent that *Mettl5*^-/-^ mice were consistently smaller than wild type (WT, +/+) and heterozygous (HET, +/-) littermates (Figure 4C). Monitoring mouse weight across several weeks revealed that this size difference persists over time in both male and female mice, with the *Mettl5*^-/-^ mice consistently weighing less than heterozygotes (Figure 4D). Though these measurements were taken from weeks 4 to 10, we noted that the difference was already present at the point of weaning (4 weeks), and persisted with similar magnitude, suggesting that this difference arose early in development, and possibly even prior to birth.

**Figure 4.**
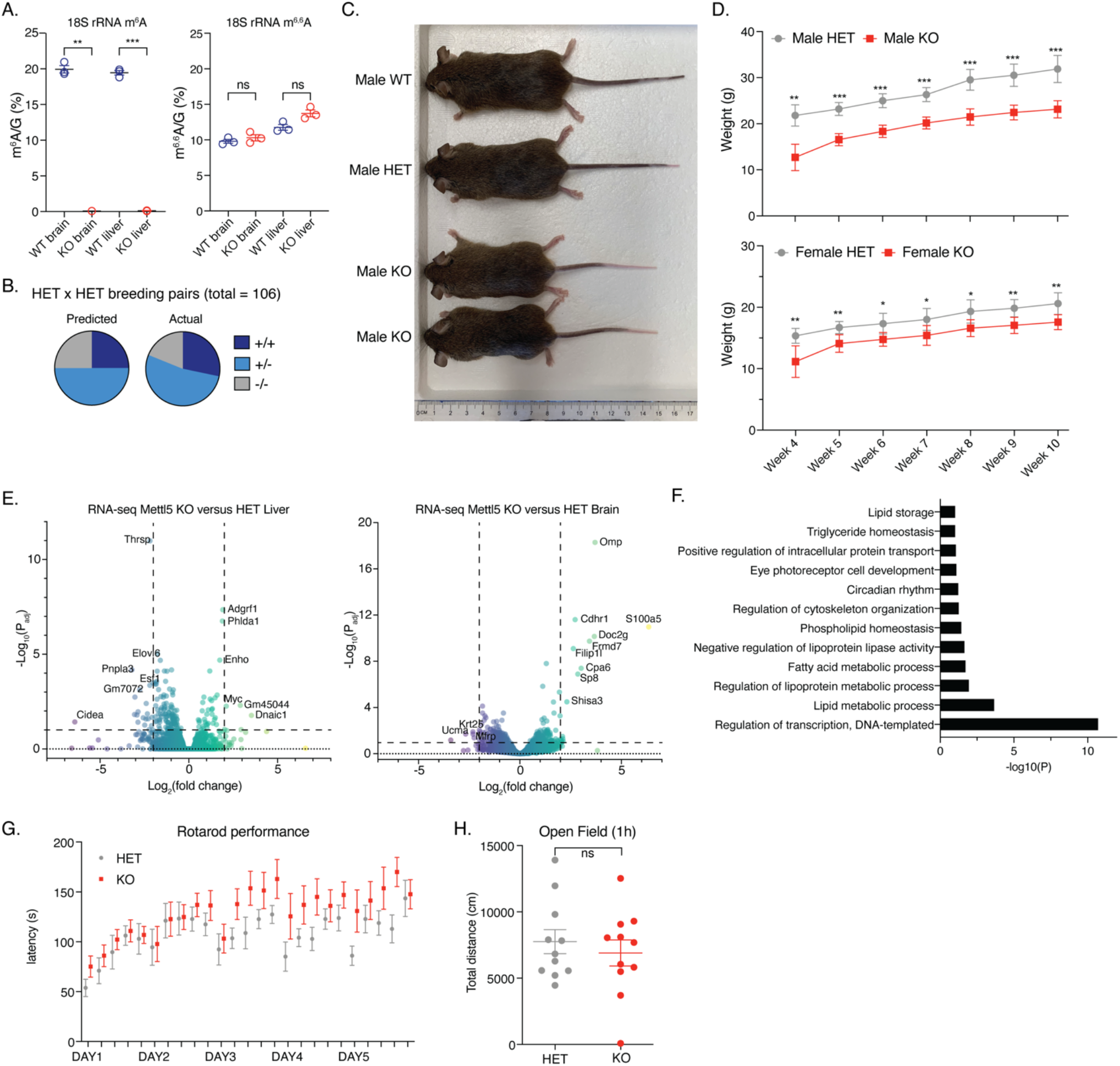
*Mettl5* knockout mice have developmental and metabolic defects. **(A)** Levels of m^6^A (left) and m^6,6^A (right), normalized to G, obtained by LC-MS/MS of 40-nt probe-purified segments of 18S rRNA surrounding the m^6^A 1832 site from brains and livers of *Mettl5*^*+/-*^ and *Mettl5*^*-/-*^ mice. n = 3 pairs, analyzed by one-way ANOVA comparison of each pair of WT and KO tissue with Sidak test for multiple comparisons. **(B)** Pie chart of the predicted (left) and actual (right) mice born of wild type (WT, +/+, dark blue), heterozygous ((HET, +/-, light blue), and knockout ((KO, -/-, grey) genotypes from heterozygous x heterozygous breeding pairs (total = 106 mice; 30 WT, 56 HET, 20 KO mice). **(C)** Photograph of male littermate WT, HET, and *Mettl5* KO mice at 8 weeks of age. **(D)** Body weights in grams of male (top) and female (bottom) HET and Mettl5 KO mice from 4 weeks (weaning) to 10 weeks of age. n=6 mice for male HET, n=4 for male KO, n=5 for female HET and KO; unpaired t-test to compare each time point. **(E)** Volcano plots from mouse liver (left) and brain (right) RNA-seq data displaying the negative log_10_ of the adjusted P-value versus the log_2_ fold change comparing the transcriptomes of *Mettl5* KO and HET mice (n = 3 pairs). **(F)** Dysregulated pathways based on analysis of RNA-seq data from *Mettl5* KO and HET mouse livers by the software DAVID. **(G)** Latency in seconds for HET and *Mettl5* KO mice to fall off of the rotarod in a rotarod performance test. **(H)** Total distance in centimeters travelled by HET and *Mettl5* KO mice when placed in a new environment in an open field test. Unpaired t-test shown. G and H: n = 11 pairs. ns: not significant, * p<0.05, ** p<0.01, *** p<0.005

In dissecting mice to harvest tissues for gene expression analysis, we also noted that *Mettl5*^-/-^ mice consistently had less body fat than *Mettl5*^+/-^ or *Mettl5*^+/+^ mice (Supp. Fig. 4E). Indeed, RNA-seq analysis of brain and liver tissues revealed gene expression patterns that suggest altered metabolism in these mice. In particular, gene ontology analysis of genes downregulated in *Mettl5*^-/-^ mouse liver revealed changes in genes involved in lipid biosynthesis and storage, consistent with the reduced body fat observed in *Mettl5*^-/-^ mice (Figure 4E,F). Noting the dramatic downregulation of *Thyroid Hormone Responsive Protein* (*Thrsp*) (Figure 4E), we measured T3 levels in blood collected from the mice used for these RNA-seq studies and found T3 levels may be elevated in the blood of *Mettl5*^-/-^ mice relative to heterozygotes (Supp. Fig. 4F).

Given the intellectual disability noted in human patients with mutations in METTL5, we also wanted to determine whether we could recapitulate any of these cognitive or behavioral changes in our mouse model. Recent reports suggest that loss of *Mettl5* results in reduced locomotor activity and exploratory activity, and defects in learning and memory (22,27). In our mouse model, we did not observe similar defects in locomotor and exploratory activity from rotarod performance and open field tests (Figure 4G,H), in fear-based learning from a shuttle box test (Supp. Fig 4G), or in instrumental learning from FR1 acquisition (Supp. Fig 4H,I). However, we note that our experiments were done comparing knockout mice with heterozygotes, not wild type mice, which combined with small variations in experimental setup could explain the different outcomes.

## DISCUSSION

In our study, we identified TRMT112 to be the primary binding partner of METTL5, a finding consistent with other recent reports (21-23). We found that protein-protein interactions between METTL5 and TRMT112 are important for METTL5 stability (Figure 1D,E; Supp. Fig. 1C,D), and methyltransferase activity (Figure 2G), though the latter is likely a consequence of the stabilizing effect of TRMT112 on METTL5. Of clinical significance, we found that the METTL5-TRMT112 interaction was severely abrogated by mutations known to cause intellectual disability in humans (Figure 1C) (18,20). Structural analysis of the three major human variants reported by Richard *et al*. and Hu *et al*. suggests that all three mutations (R115Nfs*19, K191Vfs*10, G61D) would likely disrupt proper folding (Supp. Fig. 1C). Of these, the frameshift variants, R115Nfs*19 and K191Vfs*10, were demonstrated to have lower expression (18,20). Structural analysis suggests that the METTL5^G61D^-TRMT112 complexes that do form may lack catalytic activity, since the G61D mutation creates a polar interaction with S-adenosyl methionine (Supp. Fig. 1C). Disruption of the METTL5-TRMT112 interaction likely also decreases the stability of these METTL5 variants, negatively impacting their activity and contributing to disease. To our knowledge, this is the first time that the METTL5-TRMT112 interaction in the context of human disease has been investigated. Intriguingly, our results further suggest the possibility that there are other pathways dysregulated by disease-causing METTL5. For example, the availability of TRMT112 to its other binding partners, which include AlkBH8 and HemK2, could also be dysregulated in this context (28,29,32).

We characterized METTL5 m^6^A methyltransferase activity and investigated its substrates *in vitro* and *in vivo*, identifying 18S m^6^A1832 as a major METTL5 substrate in mammalian cells, a result corroborated by recent reports (21,22,33). Intriguingly, though the nucleolar localization of METTL5 pointed to its role in rRNA methylation, we also found significant cytoplasmic localization of METTL5 by fluorescence microscopy and biochemical fractionation (Figure 2A,B). Significant cytoplasmic localization has also been reported in neurons (20) and *Drosophila melanogaster* (23), raising questions about the cytoplasmic function of METTL5. One plausible cytoplasmic role for METTL5 is late-stage methylation of rRNA, in line with an analysis by van Tran *et al*. of published ribosome structures that found density for the METTL5-TRMT112 complex near helix 44 only at late stages of processing (21,34). METTL5 may also remain associated with the ribosome in the cytoplasm, where it could regulate translation by recruiting factors to the ribosome. We note, however, that western blotting for METTL5 across polysome profiling fractions did not reveal an interaction with translating ribosomes in HeLa cells (Figure 3C). Lastly, cytoplasmic METTL5 localization could suggest the existence of other METTL5 methylation targets, and differences in abundance and localization of substrates in various cell lines could contribute to discrepancies in its localization. Our CLIP-seq and m^6^A-seq identified a small subset of transcripts that were both associated with METTL5 and whose levels changed in METTL5-KO cells, respectively, supporting this idea (Supp. Fig. 2A-D). While some of these may represent indirect interactions, it is interesting that motifs mimicking the 18S m^6^A1832 sequence context are enriched for differentially methylated m^6^A peaks on transcripts overrepresented in the METTL5 CLIP data (Supp. Fig. 2D). Moreover, there is discordance among existing studies about non-18S rRNA substrates; for instance, while Ignatova *et al*. demonstrated METTL5 activity on polyA-enriched RNA, van Tran *et al*. did not identify any non-rRNA sites in their miCLIP screen (21,22). As such, the existence and identity of other METTL5 substrates remains to be clarified through future work.

METTL5 activity on 18S rRNA prompted us to characterize the effects of METTL5 depletion on translation. Though polysome profiling did not reveal global translational changes in HeLa or HepG2 METTL5-KO cells (Figure 3B,C; Supp. Fig 3B), ribosome profiling revealed dysregulated transcripts (Figure 3E) and greater effects at the translational than the transcriptional level (Figure 3D), consistent with 18S rRNA being a major cellular substrate of METTL5. Interestingly, a more drastic effect on polysomes has been documented in mESCs (22,24,33). Clues to how METTL5 causes transcript- and context-specific effects may come from the location of the 18S m^6^A1832 site at the tip of helix 44 near the decoding center (4,21). Rong *et al*. proposed that the methyl group may fine-tune the conformation of the decoding center and suggested that the methylated adenine and its base-pairing partner are in closer proximity to mRNA in the human ribosome than in structures lacking the methylation from other organisms (24). While we and others have observed through ribosome profiling that certain transcripts and codons are more affected than others (Figure 3E, Supp. Fig 3C,E), the specific codons and transcripts affected are not consistent across datasets (22,33). Loss of 18S m^6^A1832 may also affect the position of helix 44, which is located near binding sites for key initiation and re-initiation factors, including eIF1, eIF1A, DENR, and eIF2D (35-37). Indeed, Rong *et al*. reported altered binding of the initiation factors eIF3A and eIF4E to translating ribosomes and decreased phosphorylation of the translation initiation-related signaling protein RPS6 in METTL5-KO cells (24). It is thus possible that some of the effects of METTL5 may be mediated by changes in binding of initiation-related factors to the ribosome. Considering the larger context beyond the ribosome itself may also shed light on differences in the effects of METTL5 knockout on translation in different cell lines, as there are known differences in translation-related signaling pathways among cell lines, especially in stem cells (38).

Our findings, particularly from our *Mettl5* knockout mouse model, provide insight into hereditary METTL5-related diseases caused by human variants. To our knowledge, this is the third *Mettl5* knockout mouse model reported, but the first for which 18S m^6^A loss was validated (22,27). Guided by the observations that *Mettl5*^-/-^ mice weighed less (Figure 4D) and had decreased body fat (Supp. Fig. 4E), we further investigated the metabolism of the mice through RNA-seq of their livers. We found dysregulated lipid, lipoprotein, and fatty acid metabolic pathways in knockout mice (Figure 4F) and changes in thyroid hormone signaling (Figure 4E, Supp. Fig. 4F). Interestingly, metabolic dysregulation was also suggested in a report of *metl-5* knockout in *C. elegans* (24). The mechanism(s) leading to these changes remain to be elucidated but could have clinical significance, as many patients with METTL5-associated microcephaly and intellectual disability reported by Richard *et al*. are reported to have reduced body weight (20). Unlike recent reports (22,27), however, tests with our mouse model for neurological and behavioral deficits failed to show statistically significant differences between heterozygous and knockout mice in learning ability, motor activity, or exploratory activity (Figure 4G,H; Supp. Fig. 4G,H,I), potentially due to differences in experimental details or mouse model design (Supp. Fig. 4A). Another possibility is that the loss of activity in heterozygotes may be enough to affect neurological function, although the intellectual disability and microcephaly clinical phenotypes were reported to be autosomal recessive (20). We also note that the described patient METTL5 mutants are expressed to some degree, meaning our complete knockout cell lines and mouse model do not entirely mimic the physiological conditions. It also remains possible that the neurological and behavioral defects caused by METTL5 mutations may be more subtle than is easily detectable by the simple tasks we tested but significant enough in disrupting complex tasks in humans to lead to clinical intellectual disability.

Given that 18S m^6^A1832 is currently the only validated substrate of METTL5 and the effects of METTL5 knockout seem to be most significant at the translational level, questions arise about how losing a single methyl group in the context of the whole ribosome may lead to organism-level effects. Notably, abnormalities in development and metabolism have been found in characterized clinical ribosomopathies (39). Furthermore, the differences seen in the effects of METTL5 in different cell types and tissues mirror other ribosomopathies that are very tissue-specific (9). Since our findings and those reported by others have solidified the role of METTL5 in ribosome biogenesis and function, we suggest that diseases caused by METTL5 variants could represent a previously uncharacterized ribosomopathy. This clinical relevance underscores the importance of future investigation into the mechanisms by which METTL5 mutations lead to these phenotypes. Such studies should include comprehensive identification and validation of METTL5 substrates alongside detailed investigations into changes in protein translation and signaling pathways to guide the development of clinical interventions.

## MATERIALS AND METHODS

### Cell culture

FreeStyle 293-F cells were obtained from Thermo Fisher (R79007), and HeLa (CCL-2) and HepG2 (HB-8065) cells were obtained from ATCC. HeLa and HepG2 cells were cultured in Dulbecco’s Modified Eagle Medium supplemented with 10% fetal bovine serum, sodium pyruvate, and penicillin/streptomycin (Gibco). FreeStyle 293-F cells were grown in suspension in FreeStyle 293 Expression Medium (Gibco).

### Construction and validation of knockout cell lines

HeLa and HepG2 METTL5-KO cell lines were generated via CRISPR-Cas9-mediated gene disruption. Guides (see table below) were cloned into pSpCas9(BB)-SA-Puro (PX459) V2.0, a gift from Feng Zhang (Addgene plasmid # 62988 ; http://n2t.net/addgene:62988) (41). 100 picomol of each guide and its reverse complement were phosphorylated with T4 polynucleotide kinase for 30 minutes at 37°C and then annealed by incubating at 95°C for 5 minutes and then ramping down to 25°C at 5°C per minute. Phosphorylated, annealed guides were ligated into BbsI-digested PX459 using T4 DNA ligase. Constructs were sequence verified prior to transfection into HeLa and HepG2 cells. Combinations of two constructs were used to generate deletions: PX459/METTL5-guide2 and PX459/METTL5-guide32 to generate a deletion across exon 3 and 4, and PX459/METTL5-guide2 and PX459/METTL5-guide3 to generate a deletion across exons 2, 3, and 4 (Supp. Fig. 1A). These combinations of plasmids were transfected into HeLa or HepG2 cells using Lipofectamine 2000 according to manufacturer instructions. After 48 hours, media was changed to media containing 1μg/mL of puromycin, and cells were allowed to grow for approximately a week, with media changes as needed as cell death progressed. Remaining cells were then trypsinized and diluted such that they could be plated at approximately 1 cell per well. Single cells were allowed to grow over the course of 2-4 weeks and collected and expanded as needed. METTL5 KO cell lines were identified via PCR to verify the appropriate deletion and verified by western blot.

#### Guides

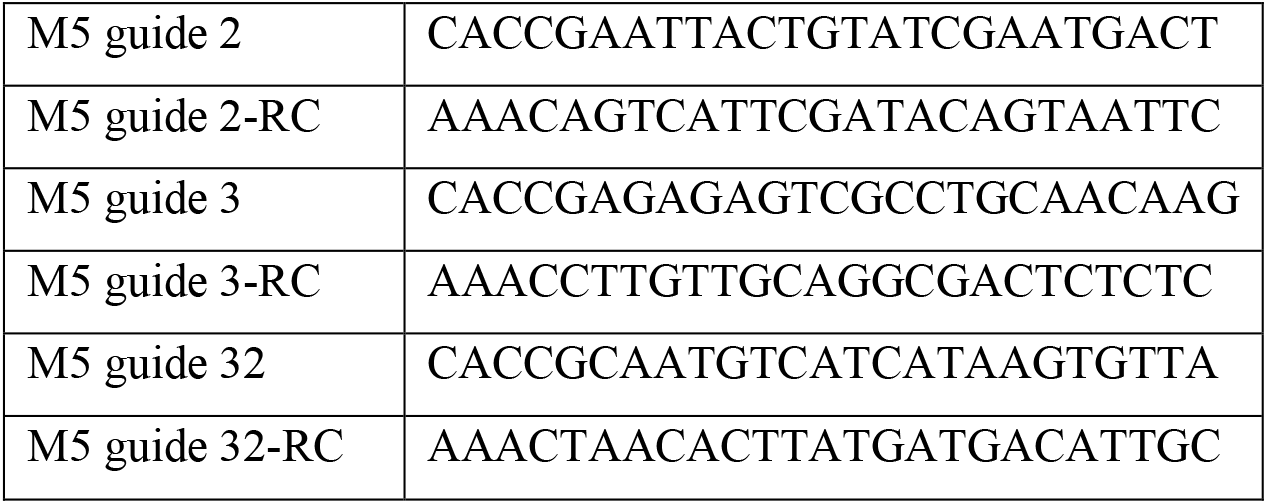

### DNA constructs

METTL5 coding sequence was amplified from pCMV-Entry-METTL5-Myc-DDK (Origene cat# PS100001) by PCR. The PCR product and pCS2-FLAG (AddGene 16331) were digested with EcoRI and XhoI and ligated with T4 DNA Ligase. Point mutations were made using quickchange mutagenesis with primers overlapping mutated sequences and encoding the following mutations: amino acids 127-129 NPP>AAA, G61D, and K191Vfs*10delAA. pCS2-FLAG was a gift from Peter Klein (Addgene plasmid # 16331; http://n2t.net/addgene:16331; RRID:Addgene 16331).

### FLAG-METTL5 co-immunoprecipitations

HeLa or Freestyle 293-F cells were transfected with constructs encoding FLAG-tagged METTL5 (WT or mutant) and were pelleted in cold 1x PBS by centrifugation at 1000g for 10 minutes at 4°C. Cells were lysed in 5 pellet volumes of lysis buffer containing 50mM Tris pH 7.4, 150mM NaCl, 10% glycerol, 1.5mM MgCl2, 0.5mM EDTA, 0.2% NP-40, 1mM DTT, and 1x SigmaFast protease inhibitor cocktail (Sigma) and rotated for 1 hour at 4°C. The lysate was clarified by centrifugation at 20,000g for 20 minutes at 4°C. For immunoprecipitation, 30μL FLAG-M2 beads per 10 million cells transfected were washed into lysis buffer 3 times before adding to clarified lysate. The mixture of lysate and beads was rotated for 1 hour at 4°C, and then washed 3 times with 0.5mL each of lysis buffer. After the final wash, 2x SDS with 100mM DTT was added to the beads and bound protein was eluted by heating at 65°C for 15 minutes, shaking at 1000rpm. Input and IP samples were separated on a 4-12% bis-tris gradient gel for western blotting (below).

### Western blots

Cells were scraped into ice cold PBS and pelleted by centrifugation at 1000g for 10 minutes at 4°C. Cell pellets were resuspended in 3 pellet volumes of lysis buffer containing 50mM Tris pH 7.4, 300mM NaCl, 1% NP-40, 0.25% deoxycholate, 1mM DTT, and 1x SigmaFast protease inhibitor cocktail (Sigma). Lysates were kept on ice for one hour with periodic agitation, and then clarified by centrifugation at 20,000g for 20 minutes at 4°C. Protein concentrations were measured by bicinchoninic acid (BCA) assay (Pierce) for normalization. Protein samples were separated on 4-12% bis-tris gradient gels using MOPS (Thermo Fisher) or MES (Thermo Fisher) buffer, and then transferred to nitrocellulose membrane by semi-dry transfer at 17V for 50 minutes using 2x NuPage Transfer Buffer (Thermo Fisher) with 10% methanol. Membranes were blocked with 5% dry milk in 1x TBST (1x TBS with 0.1% Tween-20) for 30 minutes. Primary and secondary antibodies were diluted in 5% milk in TBST and incubated as indicated in antibodies table (below). Membranes were imaged using a FluorChem R (ProteinSimple) and image analysis was performed with Fiji (42).

### RNA purification

Total RNA was isolated from cells using Trizol (Invitrogen) according to manufacturer’s instructions. After collecting cells in Trizol, 0.2mL chloroform per 1mL Trizol was added and the samples shaken vigorously for 15 seconds prior to incubating at room temperature for 2 minutes. Samples were then phase separated by centrifugation at 12,000g for 20 minutes at 4°C. The aqueous phase was collected, mixed with isopropanol (0.5mL per 1mL Trizol used), incubated at RT for 10 minutes and pelleted by centrifugation at 20,000g for 20 minutes at 4°C. The pellet was washed with 80% ethanol, centrifuged at 10,000g for 10 minutes at 4°C, and then dried and resuspended in RNase free water. Polyadenylated RNA was isolated using the Dynabeads mRNA Direct kit (Life Technologies) as per manufacturer instructions. Ribosomal RNA was depleted using the RiboMinus Eukaryote Kit v2 (Life Technologies), as per manufacturer instructions.

### siRNA transfection

One day prior to transfection, 150,000 HeLa cells were plated per 6cm cell culture dish in antibiotic free cell culture media. Media was changed to fresh antibiotic-free media just prior to transfection with 20 picomoles siRNA and Lipofectamine RNAiMAX transfection reagent (Invitrogen). 48 hours after transfection, cells were harvested either in Trizol (for qPCR analysis) or in ice cold PBS (for western blot analysis).

siRNAs used:

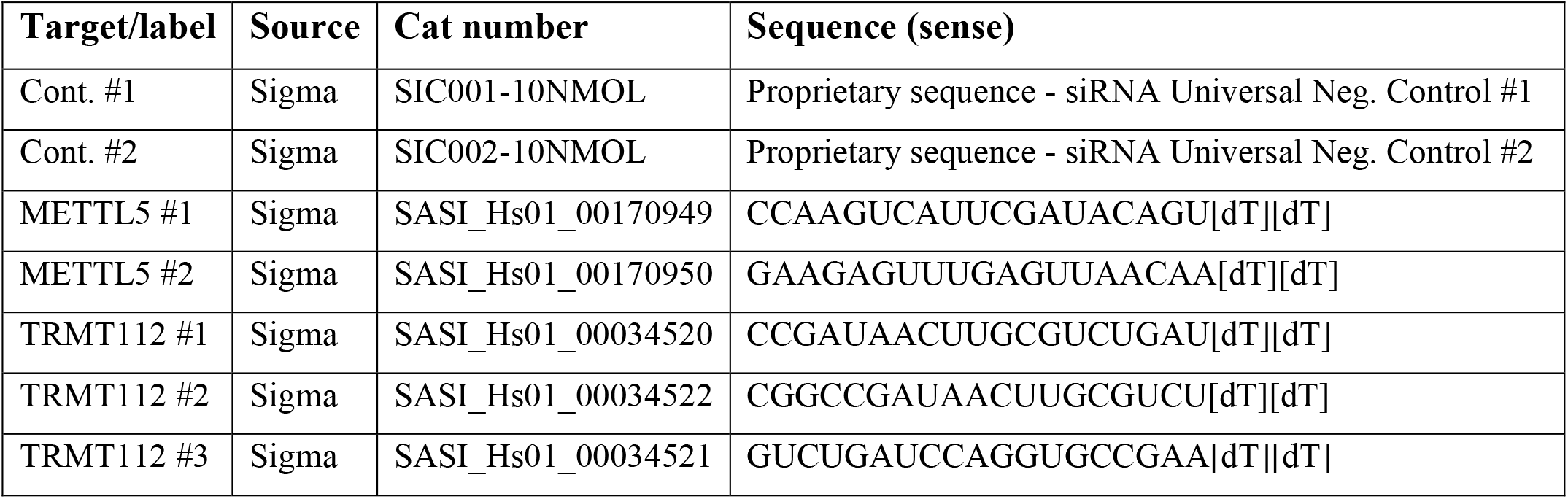

### *In vitro* methyltransferase assays

Reactions were performed by combining the desired amounts of protein and total RNA or probe in 1x methyltransferase buffer (10mM HEPES pH 7.4, 4% glycerol, 100mM KCl, 1.5mM MgCl_2_, 1mM DTT), with 2mM d_3_SAM (CDN Isotopes) and 1mM (2% v/v) SUPERaseIN RNase inhibitor (Invitrogen), for a final volume of 50μL. Reactions were incubated at 37°C for 1 hour before being snap frozen in liquid nitrogen and stored at -80°C. Prior to LC-MS/MS analysis, samples were thawed, supplemented with 1mM EDTA, and treated with Proteinase K (Sigma) at 37°C for 1 hour. Samples were purified by phenol/chloroform extraction and RNA precipitation (total RNA) or MyOne Streptavidin C1 beads (biotinylated probes; Invitrogen). For total RNA purification, an equal volume of acid phenol chloroform pH 4.5 (Invitrogen) was added to each sample and the mixture was mixed for two minutes and allowed to sit 2 minutes at RT. After centrifugation at 16,000g for 15 minutes at 4°C, the aqueous layer was transferred to a new tube. 10% the volume of 3M sodium acetate pH 5.5 and 3 volumes of 100% ethanol were added, before mixing and incubating at -80°C for 3 hours. RNA was pelleted by centrifugation at 20,000g for 20 minutes at 4°C, and then the pellets washed with 80% ethanol and resuspended in nuclease-free water. For probe purification, each sample was diluted into 350 uL of 1x IP buffer (50mM Tris pH 7.4, 300mM NaCl, 0.5mM EDTA, 1% v/v SUPERaseIN RNase inhibitor). Streptavidin C1 beads were pre-washed three times in 1x IP buffer, and 10μL bead resuspension was added to each sample. Samples were rotated for 2 hours at 4°C, washed in 1x IP buffer once, and then eluted with Trizol by shaking at RT for 20 minutes. RNA was purified from Trizol using the DirectZol RNA Miniprep kit (Zymo Research).

### LC-MS/MS

For each sample, 25-50ng of RNA was digested with 1μL Nuclease P1 (Sigma) in P1 buffer (25mM NaCl, 2.5mM ZnCl_2_) in a final reaction volume of 20μL for 2 hours at 42°C. Subsequently, 1μL of FastAP (Thermo Fisher) and 2.5μL of 10x FastAP buffer were added to each sample, and they were incubated at 37°C for 4 hours. Samples were then diluted with an equal volume of water and filtered through a 0.2μm PVDF filter (0.2μm pore size, 0.4mm diameter, Millipore). 5μL of each filtered sample was separated by reverse phase ultra-performance liquid chromatography on a ZORBAX Eclipse XDB-C18 Rapid Resolution HT 2.1×50mm, 1.8μm column (Agilent) on an Agilent Technologies 1290 Infinity II liquid chromatography system, followed by mass spectrometry on a Sciex Triple Quad 6500 triple-quadrupole mass spectrometer in positive electrospray ionization mode. Nucleosides were quantified using nucleoside-to-base transitions of 282.101>150.100 (m^6^A), 282.101>150.100 (m^1^A), 284.982>153.100 (d_3_m^6^A), 284.983>153.200 (d_3_m^1^A), 296.101>164.100 (m^6,6^A), 267.966>136.000 (A), and 284.004>152.100 (G). Standard curves were generated by injecting known concentrations of the corresponding pure nucleosides in the same run, and the percentage of modified to unmodified nucleosides were calculated based on the calibrated concentrations.

### Quantitative PCR (qPCR)

cDNA synthesis was performed in 20μL reactions consisting of 1μL random hexamer primers at 50ng/μL (Invitrogen), 1μL 10mM dNTPs (NEB), 1μL 100mM DTT (Invitrogen), 1μL SUPERaseIN inhibitor (Invitrogen), 1μL Superscript IV reverse transcriptase (Invitrogen), 4μL 5x SSIV buffer, and nuclease-free water. Samples were incubated in a thermocycler at 23°C for 10 minutes, 55°C for 10 minutes, and 80°C for 10 minutes, then held at 4°C. 10μL qPCR reactions were set up with 1μL of cDNA, 5μL FastStart Essential DNA Green Master mix (Roche), and 0.1μL each of 10μM forward and reverse primers (see table below). qPCR was performed on a LightCycler 96 (Roche) with the following settings: 95°C, 600 seconds (preincubation); 95°C 20 seconds, 60°C 20 seconds, 72°C 20 seconds (three-step amplification). Analysis was performed with LightCycler 96 SW1.1 software (Roche).

### Cell growth assays

To measure cell proliferation curves, four replicate 96-well plates were plated with 4 wells per cell line at 1000 cells per well (each plate represented one time point). For each time point, cell culture media was removed from each well and the plate was placed directly into a -80°C freezer for storage for the duration of the time course. The first time point was collected 3 hours after plating (just enough time for cells to adhere to the plate surface) and used as time 0, to normalize for small differences in plated cell number. Subsequent time points were collected at 24, 48, and 96 hours post-plating, and frozen at -80°C for at least 24 hours. The number of cells per well was quantified using the CyQuant Cell Proliferation Assay Kit (Invitrogen) as per the manufacturer’s instructions. Briefly, 200μL of lysis buffer was added to each well, pipetted up and down 4-6 times and left at room temperature for 5 minutes. 150μL of this lysate was transferred to a black, clear bottomed 96 well plate. Fluorescence was measured in a Synergy HTX plate reader (Biotek) using a 485/525 excitation/emission filter cube.

### Proteomics

Freestyle 293-F cells were seeded at 10^6^ cells/mL just prior to transfection with pCS2-FLAG-METTL5. 36 hours after transfection, cells were pelleted by centrifugation at 500g for 5 minutes at 4°C. The cells were washed with ice cold PBS and pelleted by centrifugation at 500g for 5 minutes at 4°C. The washed cell pellet was lysed in 5 pellet volumes of lysis buffer (50mM Tris pH 7.4, 50mM NaCl, 10% glycerol, 1.5mM MgCl_2_, 0.5mM CaCl_2_, 0.5% NP-40, 1x SigmaFast protease inhibitor [Sigma], 1x PhosSTOP [Sigma], and 10units/mL DNaseI [NEB]) by rotating for 1 hour at 4°C. Lysate was first clarified by centrifugation at 10,000g for 10 minutes at 4°C, and then the supernatant was transferred and clarified a second time by centrifugation at 20,000g for 20 minutes at 4°C. 500μL FLAG-M2 magnetic beads (Sigma) were washed three times with 2mL lysis buffer per wash, and added to lysate and rotated for 30 minutes at 4°C. After incubation with lysate, the beads were washed twice with 1.5mL lysis buffer, three times with 1.5mL high salt wash buffer (50mM Tris pH 7.4, 500mM KCl, 0.05% NP-40), and once with 1.5mL PBS with 0.1 Tween-20. Bound proteins were then eluted with 500uL of 2x Laemmli sample buffer (BioRad) supplemented with 100mM DTT by heating at 70°C for 15 minutes, shaking at 1000rpm. Samples were then separated on a 4-12% bis-tris gradient gel and stained by Silver Stain for Mass Spectrometry (Pierce). Bands of interest were excised using razor blades and stored in 1.5mL tubes at -80°C prior to being shipped on dry ice and analyzed by MS BioWorks, LLC (Band ID service; Ann Arbor, Michigan, USA).

### Fluorescence microscopy

20,000 cells were plated per acid-washed glass coverslip in a 24-well cell culture dish 24 hours prior to fixation with 4% paraformaldehyde in PBS for 10 minutes at RT. Coverslips were then washed 3 times with PBS and transferred to a humidity chamber. Coverslips were incubated in blocking solution (1X PBS, 1% goat serum, 10mg/mL BSA, 0.25% Triton X-100) for 30 minutes at RT, and then incubated with primary antibodies diluted in blocking buffer for 2 hours at RT. After three 5 minute washes with 1x PBS with 0.25% Triton X-100, coverslips were incubated with secondary antibodies diluted in blocking solution for 1 hour at RT. After three 5 minute washes with 1x PBS, the coverslips were mounted onto glass slides with ProLong Gold Antifade Mountant with DAPI (Life Technologies) and allowed to cure overnight at RT. Coverslips were imaged on an Olympus BX53 microscope with an Olympus U-HGLGPS light source using a 40X objective (NA 0.95). Antibodies used are listed in the table below.

### Cellular fractionation

Cellular fractionation was performed essentially as previously described (43). HeLa and HepG2 cells were scraped into ice cold PBS and pelleted by centrifugation at 500g for 5 minutes at 4°C. The cell pellet was gently resuspended and washed in 5 pellet volumes of Buffer A (10mM HEPES pH 7.5, 4mM MgCl_2_, 10mM KCl, 10% glycerol, 340mM sucrose, 1mM DTT, 1x SigmaFast Protease Inhibitor [Sigma]) three times, pelleting cells by centrifugation 500g for 5 minutes at 4°C between each wash. After the final wash, the pellet was resuspended in 2.5 pellet volumes of Buffer A. An equal volume of Buffer A + 0.2% Triton X-100 was added to this cell suspension with gentle mixing, and the samples then incubated on ice for 12 minutes with gentle inversion every 3 minutes. The lysate was then centrifuged 1200g for 5 minutes at 4°C, pelleting nuclei. The supernatant (cytoplasmic extract) was transferred to a new tube, and nuclei washed three times with Buffer A + 0.1% Triton X-100, pelleting nuclei by centrifugation at 500g for 5 minutes at 4°C between each wash and changing to fresh tubes once during these washes. The washed nuclear pellet was resuspended in one original cell pellet volume of NRB buffer (20mM HEPES pH 7.5, 75mM NaCl, 0.5mM EDTA, 50% glycerol, 1mM DTT, 1x SigmaFast Protease inhibitor [Sigma]). An equal volume of NUN buffer (20mM HEPES pH 7.5, 10mM MgCl_2_, 300mM NaCl, 0.2mM EDTA, 1M urea, 1% NP-40, 1mM DTT, 1x SigmaFast Protease inhibitor [Sigma]) was gently added to the nuclear suspension and incubated on ice for 5 minutes, with periodic inversion. After centrifugation at 1200g for 5 minutes at 4°C, the supernatant (nuclear extract) was transferred to a new tube, and chromatin pellet washed twice with a 1:1 mixture of NUN/NRB buffer with centrifugation at 5000g for 3 minutes at 4°C between each wash. The pellet was then washed twice with Buffer A in the same manner and resuspended in Buffer A after the final wash. Cytoplasmic, nuclear, and chromatin extracts were then treated with Benzonase (Sigma) for 30 minutes at 37°C before being separated by gel electrophoresis and analyzed by western blot.

### Cross-linking assisted immunoprecipitation of METTL5 sequencing

#### Immunoprecipitation

For each replicate experiment, fourteen 15cm plates of ∼70% confluent HeLa cells were transfected with 5μg of pCS2-FLAG-METTL5 plasmid per plate, using Lipofectamine LTX Reagent (Invitrogen) as per manufacturer’s instructions. 24 hours after transfection, the cells were treated with 100μM 4-thiouridine (Sigma) overnight. 36 hours after transfection, cells were washed with cold PBS and crosslinked twice with 150mJ per cm^2^ 254nM UV in a crosslinker (UVP CL-1000), with plates kept on ice for the duration. Cells were harvested by scraping in ice cold PBS and pelleting cells by centrifugation at 1000g for 10 minutes at 4°C. Cell pellets were resuspended in 5 pellet volumes of lysis buffer (50mM Tris pH 7.4, 150mM NaCl, 10% glycerol, 1.5mM MgCl_2_, 0.5mM EDTA, 0.5% NP-40, 1x SigmaFast protease inhibitor) and rotated for 1 hour at 4°C. The lysate was clarified by centrifugation at 20,000g for 20 minutes at 4°C, and an aliquot of lysate transferred to a new tube as an input sample. 200μL FLAG-M2 beads (Sigma) per 10mL lysate were washed into lysis buffer 3 times before being added to clarified lysate. The bead/lysate suspension was rotated for 2 hours at 4°C, after which the beads were washed once with 1mL lysis buffer and transferred to new tubes, then 3 times with 1mL high salt buffer (50mM Tris pH 7.4, 500mM KCl, 0.05% NP-40), followed by two more 1mL washes with lysis buffer. The beads were then washed once with 1mL Proteinase K buffer (50mM Tris pH 7.4, 75 mM NaCl, 6.25mM EDTA, 1% SDS) before resuspending the beads in 450μL of this buffer (25μL of these beads were transferred to a new tube for analysis by western blot). Proteinase K (Sigma) was pre-incubated at 37°C for 30 minutes prior to use. 50μL of Proteinase K was added to beads resuspended in Proteinase K buffer. Inputs were similarly treated with Proteinase K by adding 225μL 2x Proteinase K buffer (100mM Tris pH 7.4, 150 mM NaCl, 12.5mM EDTA, 2% SDS) and 25μL Proteinase K to 250μL of input sample. Proteinase K treatment of both input and bead samples was done by incubating samples 50°C for 1 hour, shaking at 1000rpm. 3 sample volumes of Trizol were added to each, and RNA was purified using the DirectZol kit (Zymo) as per manufacturer’s instructions. (A technical note: our efforts to perform these experiments with the RNase digestion that is usually performed to reduce background and more precisely map protein binding sites were unsuccessful (44), possibly due to the small footprint of METTL5 and/or the transient nature of interactions between RNAs and catalytic enzymes.)

#### Preparation of sequencing libraries

Sequencing libraries were prepared using the TruSeq Stranded mRNA library preparation kit (Illumina), using 30-40 ng of RNA for each input and IP sample. Samples were sequenced in one lane of an SR50 flow cell on a HiSeq4000 (50bp single-end reads).

#### Data analysis

Analysis was performed very similarly to what would be done for an RNA immunoprecipitation (IP) experiment to assess which RNA transcripts are bound to a protein of interest. Three replicate experiments were performed as described above, with corresponding input and IP samples for each replicate. Quality of .fastq files was checked using FastQC v0.11.5 (http://www.bioinformatics.babraham.ac.uk/projects/fastqc/). Adapters were trimmed using Cutadapt (45) and files were then aligned to the human genome hg38 using hisat2 v2.1.0 (46) in splice-aware mode and resulting bam files were indexed and sorted with samtools version 1.7 (47). DESeq2 (48) and R version 4.0.3 (49) were used to analyze differentially expressed transcripts, thereby identifying transcripts that were enriched in the IP samples relative to input samples.

### rRNA fragment purification

A 40-nt rRNA fragment was purified similarly to previously described (30). Briefly, 3-4μg of a biotinylated DNA probe designed to bind to the target rRNA region was combined with 33μg of total RNA in 3.33x hybridization buffer (250mM HEPES pH 7, 500mM KCl) in a total volume of 150μL. The mixture was incubated at 90°C for 7 min and then allowed to slowly cool to room temperature over 3.5 hours to allow hybridization. Single-stranded RNA and DNA were digested by adding 0.5μg RNase A (Thermo Scientific) to the mixture and incubating at 37°C for 30 minutes, then 30 units of Mung Bean Nuclease were added along with 10x Mung Bean Nuclease buffer (NEB) to a final concentration of 1x, followed by a 30 minute incubation at 37°C. Streptavidin T1 beads (20μL per sample; Invitrogen) were washed 3 times in 1mL 2.5x IP buffer (375mM NaCl, 125mM Tris pH 7.9, 0.25% NP-40), then resuspended in 100μL 2.5x IP buffer. The beads were combined with the 150μL RNA-DNA hybridization mix and rotated at 4C for 1 hour. After rotation, the beads were washed 3 times in 1x IP buffer (150mM NaCl, 50mM Tris pH 7.9, 0.1% NP-40), twice in nuclease-free water, and resuspended in 30μL nuclease-free water. The resuspended beads were heated at 70°C for 5 minute to denature the RNA from the probes, and quickly placed on ice before placing on a magnet to collect the RNA-containing supernatant.

### m^6^A-seq

#### Experiment

1μg of 1x ribo-depleted RNA per sample was fragmented using the Bioruptor Pico sonicator (Diagenode) 30 sec on, 30 sec off, 30 cycles. About 1/20^th^ of each sample was saved as input, and immunoprecipitation was performed using the EpiMark *N*^6^-methyladenosine Enrichment Kit (NEB) as per manufacturer’s instructions. Briefly, 1μL m^6^A antibody per sample was coupled to pre-washed Protein G Magnetic Beads (NEB) in 1x reaction buffer (150mM NaCl, 10mM Tris-HCl pH 7.5, 0.1% NP-40 in nuclease-free water) with orbital rotation for 30 minutes at 4°C. Beads were washed, and fragmented RNA was added in 1x reaction buffer with 2% v/v SUPERaseIN RNase inhibitor (Invitrogen) Beads were washed twice each in 1x reaction buffer, low salt reaction buffer (50mM NaCl, 10mM Tris-HCl pH 7.5, 0.1% NP-40 in nuclease-free water), and high salt reaction buffer (500mM NaCl, 10mM Tris-HCl pH 7.5, 0.1% NP-40 in nuclease-free water), then resuspended at room temperature in 30μL Buffer RLT (Qiagen) per sample. Eluates were purified using the RNA Clean & Concentrator-5 kit (Zymo Research). Libraries were then constructed using the SMARTer Stranded Total RNA-Seq Kit v2 (Takara) per manufacturer’s instructions and pooled and sequenced evenly across one lane of an Illumina NovaSeq6000 SP flow cell with 50bp paired-end reads.

#### Data analysis

Quality of .fastq files was checked using FastQC v0.11.5 (http://www.bioinformatics.babraham.ac.uk/projects/fastqc/). As instructed by the SMARTer Stranded Total RNA-Seq Kit v2 manual, the 3 bases at the start of all R2 reads were removed using Trimmomatic 0.39 (50) with the option HEADCROP:3. Adapter trimming and quality filtering were also performed with Trimmomatic using paired-end mode with options LEADING:3, TRAILING:3, SLIDINGWINDOW: 4:15, and MINLEN: 21. To check for contamination, files were aligned to the mycoplasma genome using hisat2 v2.1.0 (46) and any matching reads were discarded. Files were then aligned to the human genome hg38 using hisat2 v2.1.0 in splice-aware paired-end mode and resulting bam files were sorted with samtools version 1.7 (47). Using the MeRIPTools package (51), reads were counted with the function CountReads, and peaks were called using Fisher’s exact test with the function callPeakFisher. The R package QNB was used for inferential testing and differentially methylated peaks were called with an adjusted P-value < 0.1 (52). Motif searches were performed using HOMER v4.11 (53). For a background reference, sequences were extracted from random 200bp peaks that were sampled from an mRNA transcript (51).

### Polysome and ribosome profiling

Polysome profiling was performed similarly to previous reports (54,55). Four 15cm plates per sample were cultured to ∼80% confluence. Each plate was treated with cycloheximide at a final concentration of 100μg/mL in antibiotic-free medium and incubated at 37°C for 7 minutes. Plates were washed with 10mL of ice-cold PBS with 100μg/mL cycloheximide twice, then cells were collected in ice-cold PBS and centrifuged at 500g for 5 minutes. Each pellet was resuspended in 3 pellet volumes of lysis buffer (20mM HEPES pH 7.6, 100mM KCl, 5mM MgCl2, 1% Triton x-100, 100μg/mL cycloheximide, 1% v/v SUPERaseIN inhibitor in nuclease-free water) and cells were lysed with rotation at 4°C for 20 minutes before being centrifuged at 16,000g for 15 minutes. To the clarified lysate, 4μL of Turbo DNase (Thermo Fisher) was added and samples were incubated at room temperature 15 minutes then centrifuged again to clear the lysate. The absorbance at 260nm of each lysate was measured by Nanodrop (Thermo Fisher) and samples were adjusted to the same optical density with lysis buffer. One fifth of each sample was saved as input, while the remaining portion of each sample was loaded on a 5-50% w/v sucrose gradient prepared in lysis buffer in a SETON Scientific open top polyclear tube (cat# 7042). Samples were then centrifuged at 28,000 rpm for 3 hours at 4°C using an Optima L-100XP centrifuge with an SW28 rotor. After centrifugation, absorbance was read and fractions were collected by Gradient Station (BioComp) equipped with an ECONO UV monitor (BioRad) and fraction collector (Gilson). The max absorbance (AUFS) was set to 0.5, number of fractions to 30, and distance/fraction to 3.2mm.

Ribosome profiling was performed similarly, with a few key differences. Firstly, after DNase treatment, about 20% of the sample was saved as input with 3 volumes of Trizol added, while the other 80% was treated with 3μL MNase (NEB) in 1x MNase buffer for 15 minutes at room temperature to digest RNA not covered by ribosomes. Additional SUPERaseIN (Invitrogen) was added to a final concentration of 0.5mM to quench the reaction. Then, samples were loaded on 5-50% w/v sucrose gradients and fractionated as above. Secondly, after fractions were collected, samples were processed for sequencing. Fractions containing the digested monosomes were collected and pooled, and three volumes of Trizol (Invitrogen) was added to each sample. RNA purification was performed for both input and ribosome-protected fragment (RPF) samples by chloroform extraction followed by isopropanol precipitation, and one round of rRNA depletion was performed (see “RNA purification” section). Input RNA was fragmented by combining 20μL RNA with 7.5μL PNK buffer A (Thermo Fisher) and incubating at 94°C for 25 minutes, then end repaired by diluting in 37.5μL nuclease-free water, adding 10μL T4 PNK (Thermo Fisher) and incubating at 37°C for 30 minutes. Then, 12μL 10mM ATP (EMD Millipore), 2μL 10x PNK buffer A, and 6μL T4 PNK were added to incubate at 37°C for 30 minutes. These samples were purified using the RNA clean and concentrator-5 kit (Zymo Research). Purified RPF RNA was run in 1x Novex TBU sample buffer (Thermo Fisher) on a 10% TBE-urea gel (Thermo Fisher) in 1x RNase-free TBE buffer (Thermo Fisher) with an IDT 10/60 loading control (IDT) for 1 hour at 180V. The gel was stained 15 minutes in 1x SYBR gold in TBE with gentle shaking and visualized using a Gel Doc EZ Imager (Biorad). Fragments between 26-34 nucleotides were excised and purified by Small-RNA PAGE recovery kit (Zymo Research) per manufacturer’s instructions. RPFs were end repaired similarly to inputs with T4 PNK. Input and RPF RNA samples were used to prepare libraries for sequencing using the NEBNext Small RNA Library Prep Kit (NEB) per manufacturer’s instructions, and were sequenced by Illumina NovaSeq6000 with 100bp single-end reads on two SP flowcells.

#### Data analysis

Analysis of polysome profiles was performed using Microsoft Excel and GraphPad Prism. For ribosome profiling, fastq files from the two flowcells were merged using the command “cat” and data quality was checked with FastQC v0.11.5 (http://www.bioinformatics.babraham.ac.uk/projects/fastqc/). Adapters were trimmed and quality filtering was performed with BBDuk from BBTools version 38.84 using a reference fasta file of the NEBNext adapter sequences and settings ktrim = 4, qtrim = rl, trimq = 20, k = 31, mink = 11, hdist = 1, entropy = 0.5, entropywindow = 30, entropyk = 5, minlen = 25 and maxlen = 34 (for RPFs), and minlen = 15 (for inputs) (BBtools: (https://sourceforge.net/projects/bbmap/). For input files, alignment to human genome hg38 (56) was performed using STAR 2.7.3a (57) in quantMode TranscriptomeSAM. Bam files were sorted and uniquely mapped reads extracted with samtools v1.7 (47), followed by featureCounts v1.6.0 (58) to create a counts table from mapped reads with options - s 0, -g gene_id, and -t exon. rRNA reads were removed from RPF Fasta files using BBDuk with a reference fasta file of human rRNA sequences. Analysis was then performed on the RiboToolkit server (http://rnabioinfor.tch.harvard.edu/RiboToolkit/) (40). First, RPF files were uploaded for single-case submission in collapsed fasta format with the following options: species = homo sapiens (hg38), RPF interval = 25-34, allowed mismatch = 2, max multiple-mapping = 1, no duplicate removal, offsets to infer P-sites - calculate by RiboToolkit, min coverage for pause sites = 10, fold change for pause sites = 20, ORF p-value = 0.05. Input counts table and single case Job IDs were then used as inputs for group case analysis with F value = 2 and P-value = 0.05.

### RNA-seq, METTL5-KO HeLa cells

#### Experiment

3 biological replicates of wild type and 2 different clones METTL5-KO HeLa cells (clone 1 and 7) were grown to confluency prior to lysis in Trizol (Invitrogen). Total RNA was purified as described above (‘RNA purification’) and sequencing libraries were generated using an HIV reverse transcriptase evolved for m^1^A detection, as previously described (59). Libraries were sequenced using an Illumina NovaSeq6000 on an S1 flowcell with 100bp paired end reads. For RNA-seq analysis, only read 1 (R1) was used.

#### Analysis

Quality of fastq files was checked using FastQC v0.11.5 (http://www.bioinformatics.babraham.ac.uk/projects/fastqc/). Adapters were trimmed using Cutadapt (45). To check for contamination, files were aligned to the mycoplasma genome using hisat2 v2.1.0 (46) and any matching reads were discarded. Files were then aligned to the human genome hg19 (56) using hisat2 v2.1.0 (46) and resulting bam files were indexed and sorted with samtools version 1.7 (47). Differential expression analysis was performed using DESeq2 (48) and R version 4.0.3 (49). Gene ontology analysis was performed with MetaScape (60).

### RNA-seq, mouse tissues

#### Experiment

Mouse organs were collected immediately upon sacrifice, washed in ice-cold PBS, placed in ∼4 volumes of Trizol (Invitrogen), and then sonicated using a hand-held sonicator (OMNI International) with a few brief pulses until there were no more visible chunks. Samples were stored at -80°C until use. Libraries were prepared using the SMARTer Stranded Total RNA-Seq Kit v2 (Takara) per manufacturer’s instructions and sequenced on a NovaSeq6000 with 100 bp single-end reads.

#### Data analysis

Sequencing data quality was checked by FastQC v0.11.5 (http://www.bioinformatics.babraham.ac.uk/projects/fastqc/). Adapter trimming and quality filtering was performed with Trimmomatic (50) in single-end mode with options LEADING:3, TRAILING:3, SLIDINGWINDOW:4:15, and MINLEN:36. Reads were then aligned to the mm10 genome (56) using STAR 2.7.3a (57). Read counts mapping to each gene were obtained using FeatureCounts (58) with options -s 2, -g gene_id, and -t exon. Differential expression analysis was then performed in R version 4.0.3 (49) using DESeq2 1.28.1 (48). Gene ontology analysis was performed with MetaScape (60).

### Knockout mouse generation and mouse husbandry

*Mettl5* knockout mice were generated at the Gene Targeting & Transgenic Facility at Janelia Research Campus. A *Mettl5* conditional knockout line was made in which exon 2 was floxed, such that presence of Cre would induce exon 2 deletion by creating a frame shift mutation. The construct was electroporated with gRNA/Cas9 protein into ES cells that are F1 hybrid of C57bl/6 × 129S6. F1 mice transferred from Janelia Research Campus to the University of Chicago where the second generation was backcrossed to the B6 strain. To generate a whole body knockout, the conditional knockout strain described above was then crossed with the B6.C-Tg(CMV-cre)1Cgn/J line (The Jackson Laboratory, 006054) (61). All mice were housed in a specific pathogen-free facility. Both male and female mice were used throughout the study, between 4 weeks and 6 months of age. All experiments were approved by the University of Chicago Institutional Animal Care and Use Committee.

### ELISA

Blood was collected retro-orbitally from mice upon sacrifice, allowed to coagulate at room temperature for about one hour, and centrifuged at 10,000g for 15 minutes at 4°C. The supernatant (serum) was transferred to a new tube and stored at -80°C. The thyroid hormone (T3) concentration was assayed using the Biomatik Mouse Tri-iodothyronine, T3 ELISA Kit (Cat# EKC40189), per manufacturer’s instructions.

### Mouse behavioral assays

#### Rotarod Experiment

A computer-controlled rotarod apparatus (Rotamex-5, Columbus Instruments) with a rat rod (7cm diameter) was set to accelerate from 4 to 40 revolutions per minute over 300 seconds, and time to fall was recorded. Mice received five consecutive trials per session, one session per day (about 30 seconds between trials).

#### Open field

Open field chambers were 40cm x 40cm (Med Associates) with lighting at 21 lux. Infrared beams recorded the animals’ locomotor activity and rearing movements (vertical activity). Mice were put in the open filed chamber for 1 hour to record their activity.

#### Passive avoidance

The shuttle box used contained two chambers: one chamber illuminated and the other dark (Kinder Scientific). Mice were transported to the behavior room and were handled for three minutes for 3 days before the passive avoidance experiment. During tasks, the right chamber remained illuminated while the left chamber remained dark. Training began by placing the mouse into the illuminated chamber facing away from the shut guillotine door. The mouse was allowed to explore the illuminated chamber for 2 minutes. The door was then opened to let the mouse explore both the illuminated and dark chambers for 5 minutes. At the end of this exploration period, the door was shut after returning the mouse into the illuminated chamber. Two minutes later, the door was opened. Latency to step into dark chamber was recorded by the computer as the baseline. Upon entering the dark chamber, the door was closed and one foot shock (0.2 mA, 2 seconds) was delivered. Ten seconds later, the mouse was removed from the dark chamber and put back to the home cage. After 24 hours, the mouse was put into the light chamber for 2 minutes and then the latency to step into dark chamber was recorded as the 24 hour memory.

#### FR1

Mice were food deprived each day before the experiment. Mice had unlimited food access for 3 hours each day after the experiment. Each mouse was first trained in an illuminated operant box (Med Associates, St. Albans, VT, USA) on a FR1 schedule for 30 minutes each day for about three weeks, with additional random delivery of a food pellet for the first 15 minutes (one pellet per minute on average). The maximum number of food pellets by pressing the lever was set at 40. Mice were then trained on a FR1 schedule without free delivery of the food pellets. After the mice achieved more than 15 presses for two consecutive days, the results of the last of day FR1 were compared between genotypes.

### Statistical analyses

Statistical analyses were done in GraphPad Prism. Throughout the figures, replicates are plotted as individual points, along with mean and standard error of the mean (s.e.m.). For experiments comparing two conditions, unpaired t-test was used. For experiments with multiple comparisons, one-way ANOVA was used to test specific comparisons as indicated (that is to say, not every single sample was compared to every other sample). A Dunnett’s test to correct for multiple comparisons when all comparisons were made to a single control (Figures 1F, 2E; Supp. Fig. 2G). When multiple comparisons were made between experimentally determined pairs, the Sidak method was used to correct for multiple comparisons (Figure 4A, Supp. Fig. 4G). In experiments with multiple variables and parameters tested, a two-way ANOVA with the appropriate multiple comparison correction was used (Sidak test for Figure 2F, Tukey test for Figure 2G).

### Table of primers and oligonucleotides

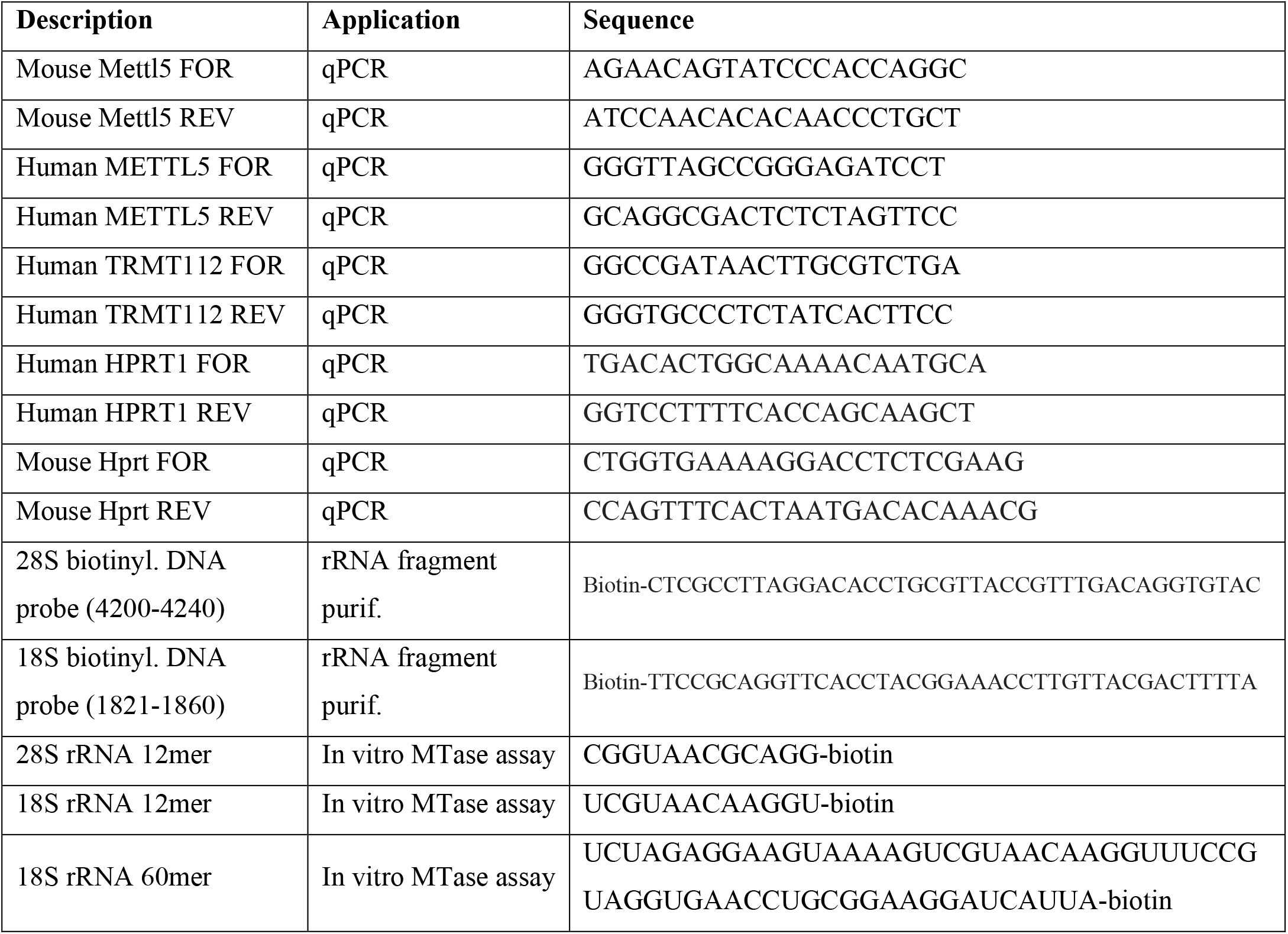

### Antibodies

| **Target** | **Source/catalog number** | **Application/species/dilution** |
| --- | --- | --- |
| anti-METTL5 | ProteinTech/16791-1-AP | Western blot/rabbit/1:500 dilution |
| anti-METTL5 | Atlas/SAB2101471 | IF/rabbit/1:250 dilution |
| anti-TRMT112 | Santa Cruz Biotechnology/sc-398481 | Western blot/mouse/1:100 dilution |
| anti-FLAG-M2 | Sigma/A8592 | Western blot/mouse/1:10,000 dilution |
| anti-RPL4 | Proteintech/11302-1-AP | Western blot/rabbit/1:1000 dilution |
| anti-RPS6 | Abcam/70227 | Western blot/rabbit/1:1000 dilution |
| anti-EIF3A | Cell Signaling Technology/3411 | Western blot/rabbit/1:1000 dilution |
| anti-EIF4E | Cell Signaling Technology/2067 | Western blot/rabbit/1:1000 dilution |
| anti-GAPDH-HRP | GenScript/A00192-100 | Western blot/goat/1:10,000 dilution |
| anti-SNRP70 | Abcam/ab83306 | Western blot/rabbit/1:1000 dilution |
| anti-histone H3 | Cell Signaling Technology/D1H2 | Western blot/rabbit/1:2000 dilution |
| anti-fibrillarin | Abcam/ab4566 | IF/mouse/1:250 dilution |
| goat-anti-rabbit-488 | Thermo Fisher/A11008 | IF secondary, 1:500 dilution |
| goat-anti-mouse-594 | Thermo Fisher/A11005 | IF secondary, 1:500 dilution |
| Goat anti-Rabbit HRP | Thermo Fisher/A16110 | Western blot, secondary, 1:5000 dilution |
| Goat anti-Mouse HRP | Thermo Fisher/A16078 | Western blot, secondary, 1:5000 dilution |

## DATA AVAILABILITY

Raw and processed data files from all high throughput sequencing experiments have been deposited in the NCBI Gene Expression Omnibus (GEO) with the following accession numbers: GSE174435 (METTL5 CLIP-seq), GSE174503 (RNA-seq of HeLa cells), GSE174420 (m^6^A-seq), GSE174418 (RNA-seq of mouse tissues), GSE174419 (ribosome profiling).

## ACKNOWLEDGEMENTS

We thank Gregory Sepich-Poore (UC San Diego) and Shun Liu (Uchicago) for bioinformatics support, Alex Ruthenburg (Uchicago), Lisheng Zhang (Uchicago), Hailing Shi (Broad Institute), and Honghui Ma (Tongji University) for advice on experiments, Richard Jones (MS BioWorks) for proteomics guidance, and Yawei Gao (Tongji University) for guidance on mouse model development. We thank Caiying Gao and the staff at the Gene Targeting & Transgenic Facility at Janelia Research Campus at HHMI for the initial phases of mouse model development, and the University of Chicago Animal Resources Center for housing and caring for the mice. Sanger and NGS sequencing were performed by the University of Chicago Comprehensive Cancer Center Sequencing Facility and the University of Chicago Genomics Facility, respectively. The work was supported by the National Institutes of Health NHGRI RM1 HG008935 to C.H. and R01DA043361 to X.Z.. C.S. and A.C.Z were supported by NIH Medical Scientist Training Program grant T32GM007281 and by NCI F30 CA253987 (C.S.) and NCI F30 CA247175 (A.C.Z.). C.H. is an investigator of the Howard Hughes Medical Institute (HHMI). S.N. is the William Raveis Charitable Fund Breakthrough Scientist of the Damon Runyon-Dale F. Frey Award (DFS-34-19) and was previously an HHMI Fellow of the Damon Runyon Cancer Research Foundation (DRG-2215-15).

## Supporting information

### Supplementary information associated with this manuscript

Supplementary figures 1-4

Table S1. Proteomics analysis of METTL5 binding partners

Table S2. Analysis of transcripts isolated from FLAG-METTL5 CLIP

Table S3. METTL5 knockout versus wild type HeLa m6A-seq differentially methylated peaks Table S4. Differentially expressed genes in HeLa-METTL5-KO cells relative to HeLa-WT cells Table S5. METTL5 knockout versus wild type HepG2 ribosome profiling data

Table S6. METTL5 knockout versus wild type mouse RNA-seq data

**Supplementary Figure 1.**
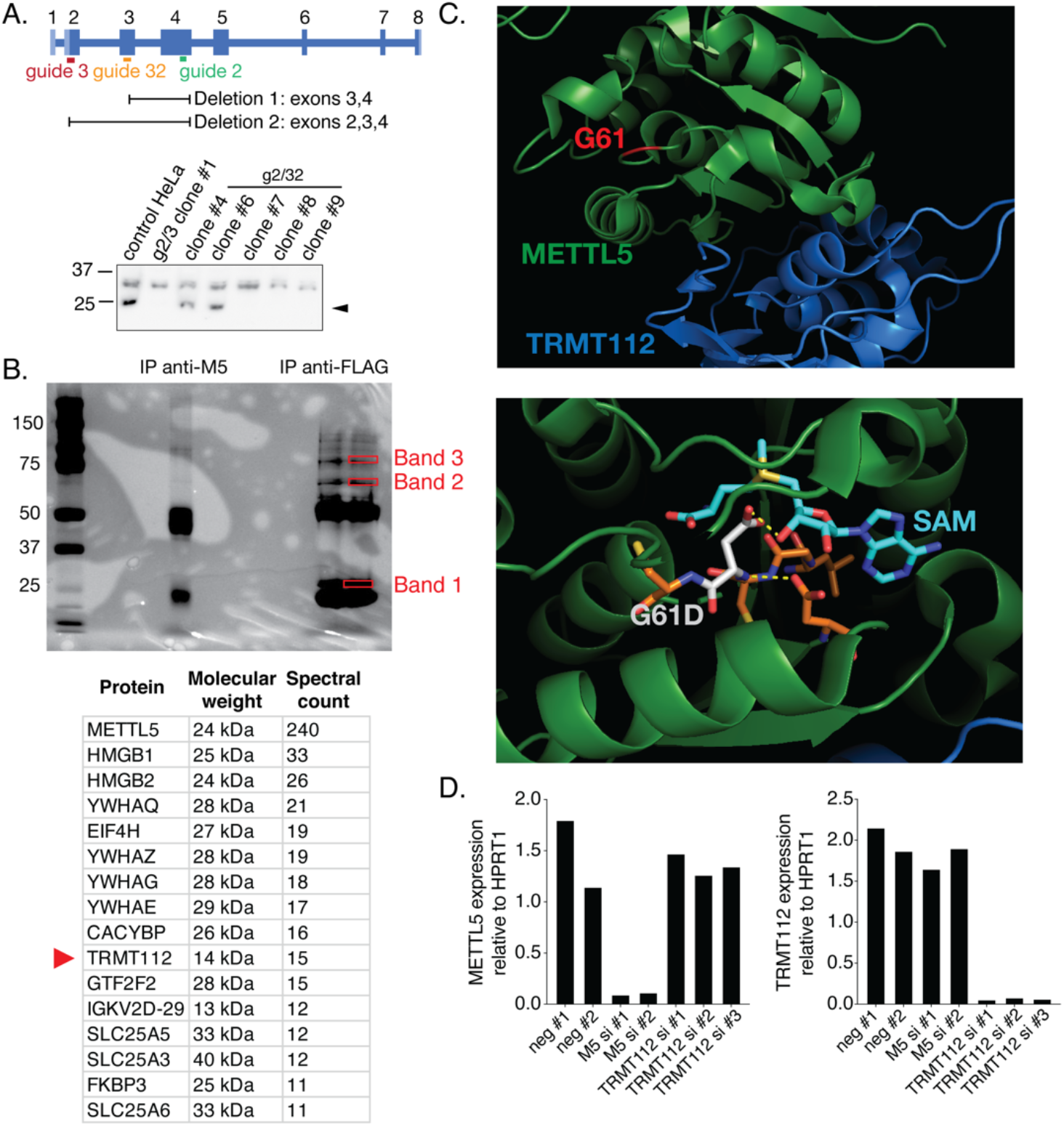
Identification and investigation of METTL5-TRMT112 interactions. **(A)** Top: schematic of human METTL5 locus showing target locations of guide RNAs used to make METTL5 knockout HeLa and HepG2 cells in this study. Guides 2 and 3 were used to generate HeLa clone 1, while guides 2 and 32 were used to generate all other HeLa and HepG2 clones in this study. Bottom: METTL5 expression in cell lines as evaluated by western blot (top) with anti-GAPDH loading control (bottom). Arrowhead signifies METTL5 band. **(B)** Top: Silver-stained polyacrylamide protein gel with protein size marker, immunoprecipitate from pulldown with endogenous METTL5 antibody, and immunoprecipitate from pulldown with anti-FLAG antibody in Freestyle 293-F cells overexpressing FLAG-tagged METTL5. Boxed bands were cut out for proteomics analysis. Bottom: Table of top hits from proteomics analysis of band 1 with molecular weight and spectral count of each. Suspected common contaminant proteins were removed from the list, but complete information, as well as analysis of bands 2 and 3, is available in Supplementary table 1. **(C)** G61D human variant mutation site displayed on the METTL5-TRMT112 structure from van Tran *et al*. (PDB: 6H2U (1)). Top: Position of G61 highlighted in an unstructured loop (red) in the context of the complex with METTL5 (green) and TRMT112 (blue). Bottom: Interactions between the mutated G>D residue (gray) and S-adenosyl-methionine (cyan). Neighboring residues are highlighted in orange. Images created with PyMOL v2.4.0 (2). **(D)** METTL5 (top) and TRMT112 (bottom) expression relative to HPRT1 (housekeeping gene) in total RNA purified from HeLa cells treated with negative control siRNAs or siRNAs targeting METTL5 or TRMT112, as evaluated by quantitative PCR.

**Supplementary Figure 2.**
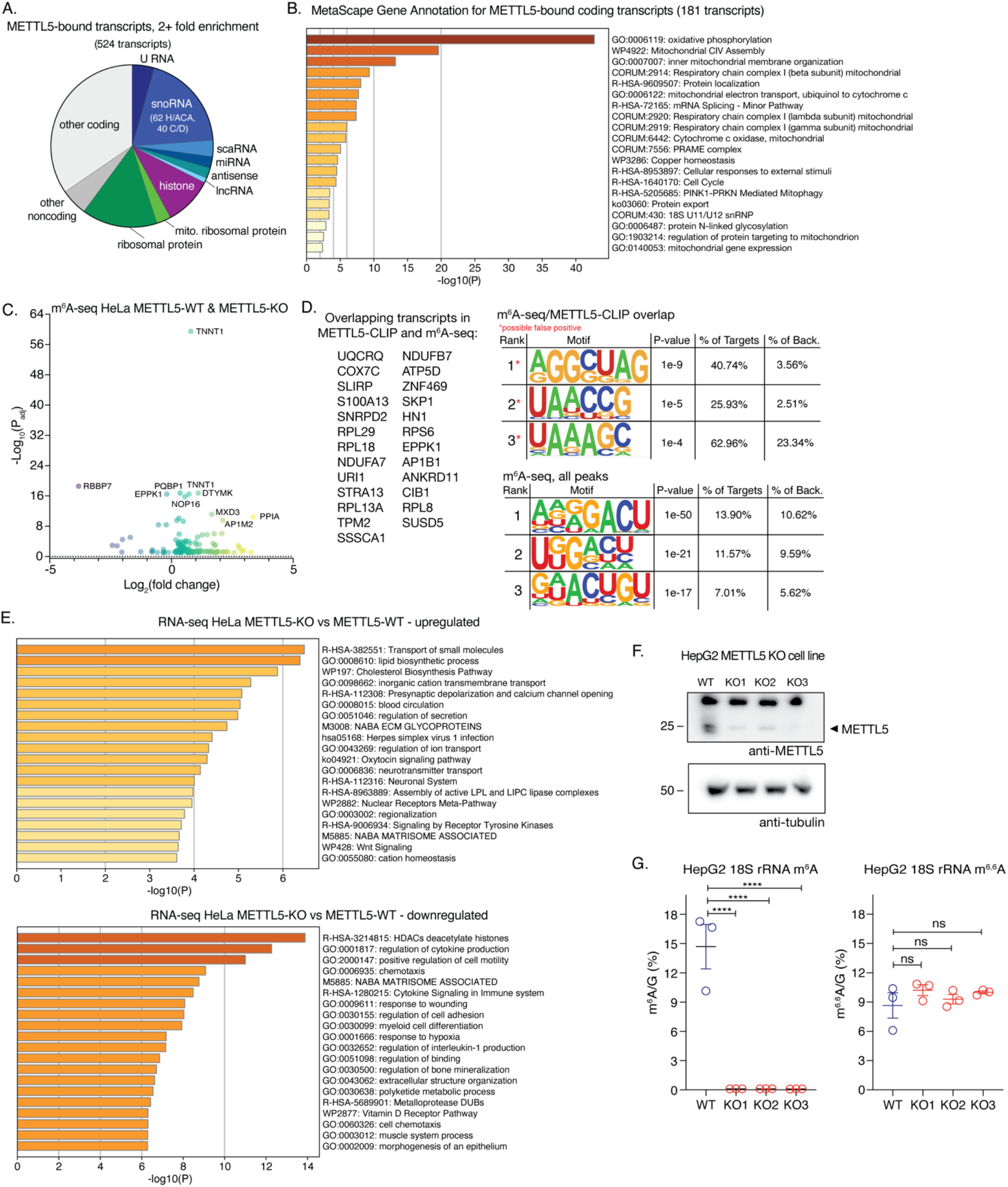
Investigation of METTL5 substrates in human cells. **(A)** Pie chart of METTL5-bound transcripts with ≥2-fold enrichment in METTL5 immunoprecipitate after crosslinking compared to input. **(B)** MetaScape gene annotations for METTL5-bound coding transcripts. **(C)** Volcano plot of differentially methylated m^6^A peaks between HeLa METTL5-KO and METTL5-WT cells, shown comparing negative log_10_ of the adjusted P-value with log_2_ fold change. **(D)** Left: List of transcripts with both differentially methylated m^6^A peaks from Me-RIP-seq and ≥2-fold enrichment in METTL5 cross-linking and immunoprecipitation sequencing in HeLa cells. Right: Common motifs in this list of overlapping targets (top) and in all m^6^A peaks from Me-RIP-seq (bottom), as predicted by HOMER (3). Back.: background (see Materials and Methods) **(E)** MetaScape gene annotation terms from transcriptionally upregulated (top) and downregulated (bottom) transcripts in RNA-seq of METTL5 knockout (KO) versus wild type (WT) HeLa cells. **(F)** Western blot analysis of METTL5 levels in wild type (WT) and knockout (KO) HepG2 samples, each expanded from a single isolated clone (top) with anti-tubulin loading control (bottom). **(G)** Levels of m^6^A (left) and m^6,6^A (right), normalized to G, obtained by LC-MS/MS of 40-nt probe-purified segments of 18S rRNA surrounding the m^6^A 1832 site from wild type and METTL5 knockout HepG2. Analyzed by one-way ANOVA, comparing all samples to WT, with Dunnett’s test for multiple comparisons. ns: not significant, * p<0.05, ** p<0.01, *** p<0.005, **** p<0.0001. Panels B and E modified from MetaScape output (4).

**Supplementary Figure 3.**
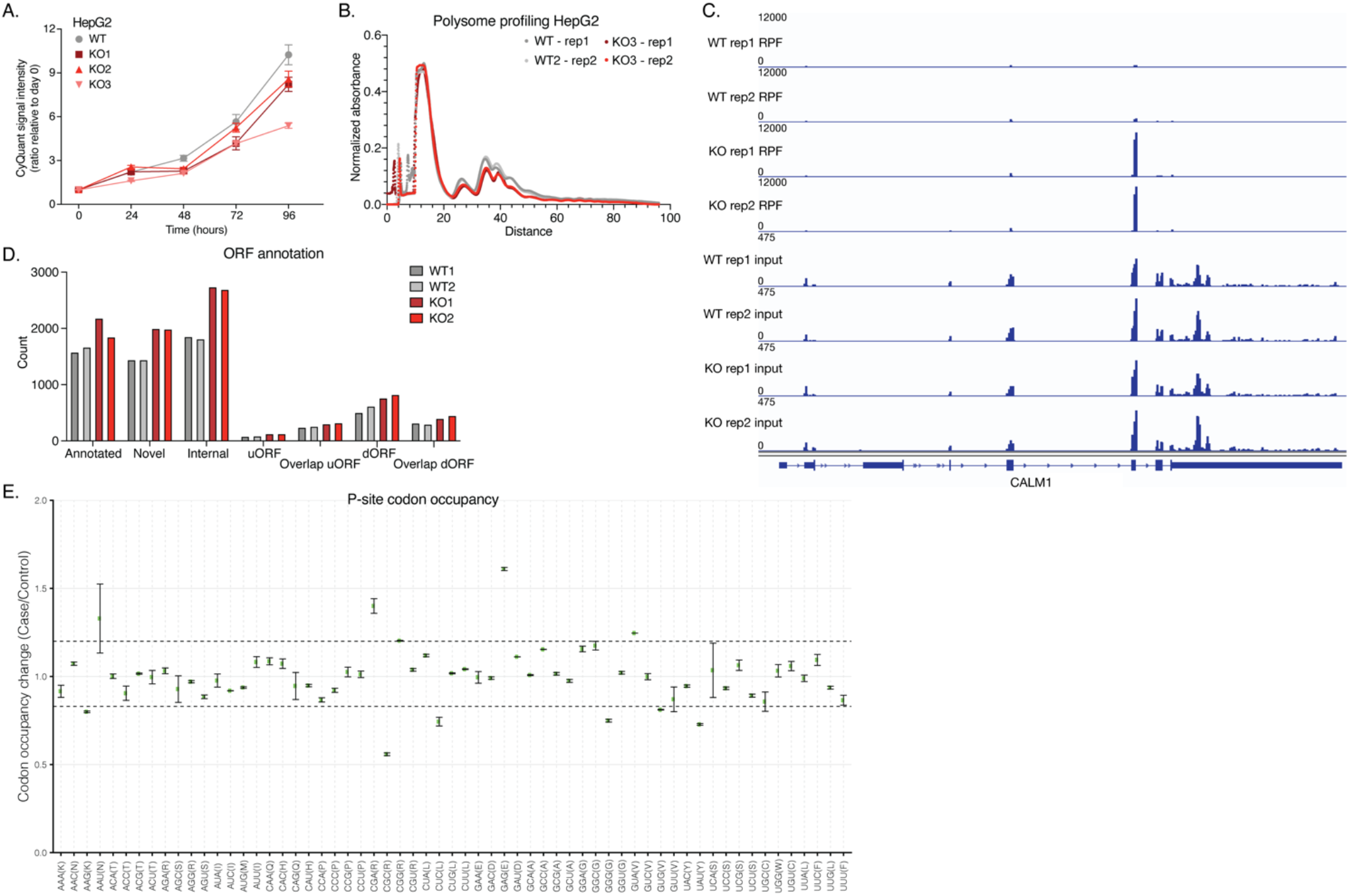
The effects of METTL5 knockout on translation and cell state. **(A)** Cell proliferation of HepG2 METTL5-WT and METTL5-KO cell lines over time as measured by CyQuant assay. Means (squares) and SDs (bars) are indicated for 4 replicate wells per condition. **(B)** Polysome profiles from HepG2 METTL5-WT and METTL5-KO cells as measured by normalized absorbance over a 5-50% sucrose gradient. **(C)** Visualization of reads at the *CALM1* locus from input and ribosome-protected fragment (RPF) samples from HepG2 METTL5-WT and METTL5-KO cells, adapted from IGV. **(D)** Active open reading frame comparison statistics of RPF sequencing read counts from METTL5-WT and METTL5-KO HepG2 cells. **(E)** Codon occupancy changes at the P site between METTL5-KO (case) and METTL5-WT (control) HepG2 cells. D and E are modified from output of RiboToolKit (5).

**Supplementary Figure 4.**
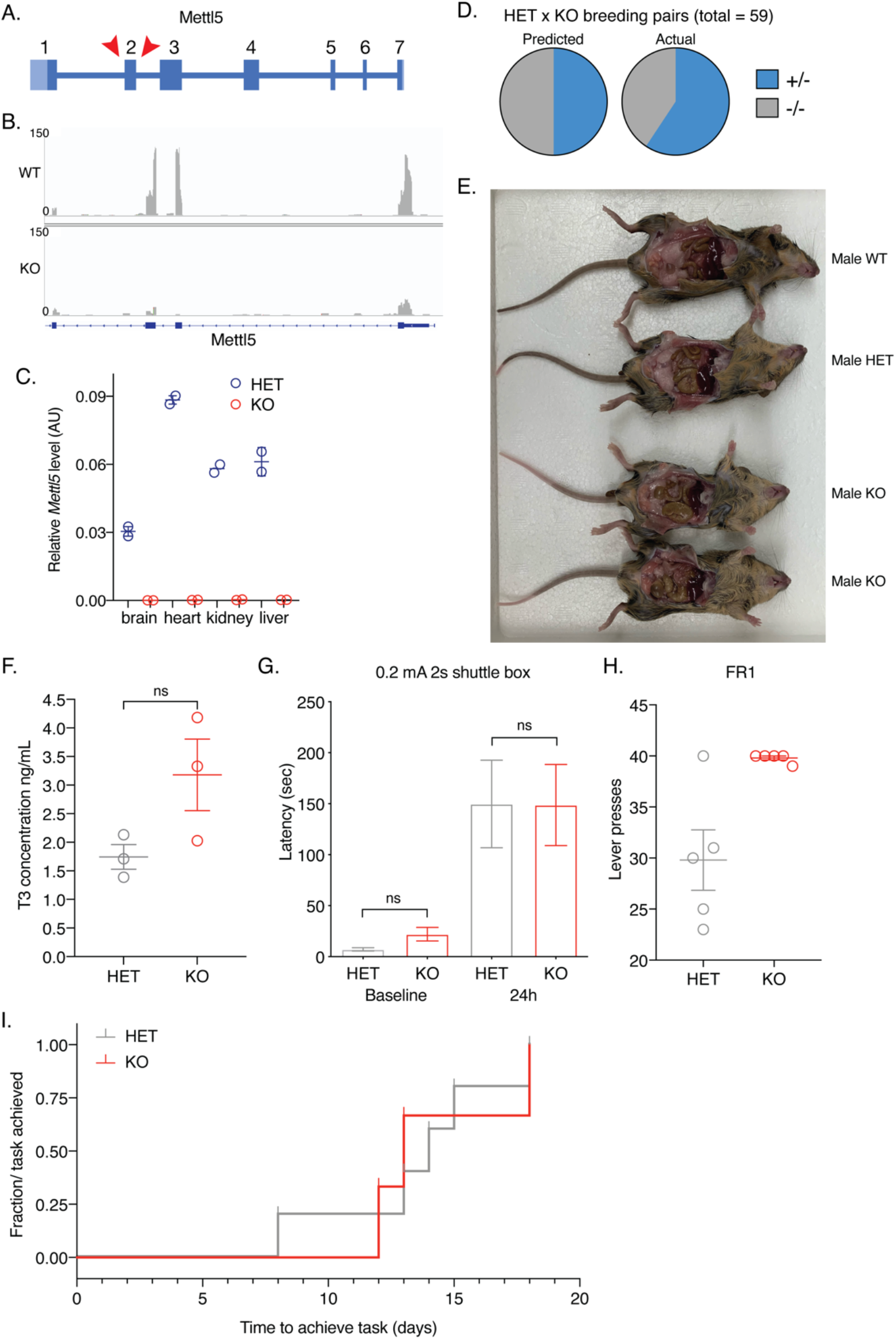
Phenotypic characterizations of *Mettl5*^*-/-*^ mice. **(A)** Schematic of mouse *Mettl5* locus showing region targeted for removal by CRISPR-Cas9 to create KO mice for this study. **(B)** Visualization of RNA-seq reads at mouse *Mettl5* locus from a littermate pair of one *Mettl5*^*+/-*^ and one *Mettl5*^*-/-*^ mouse, adapted from IGV. Exon 1 is at far right and exon 4 is on the far left; partial view shown since *Ssb* overlaps with *Mettl5*. **(C)** Relative *Mettl5* transcript level (normalized to *Hprt*) as measured by qPCR of total RNA from brains, hearts, kidneys, and livers of *Mettl5*^*+/-*^ and *Mettl5*^*-/-*^ mice. **(D)** Pie charts of the predicted (left) and actual (right) mice born of HET (+/-, light blue) and KO (-/-, grey) genotypes from HET x KO breeding pairs (total = 59 mice; 35 heterozygous, 24 knockout). **(E)** Photo of dissected male littermate mice at 8 weeks showing abdominal fat content. **(F)** T3 concentration (ng/mL) from enzyme-linked immunosorbent assay (ELISA) of thyroid hormone in serum of the same heterozygous and knockout mice used for RNA-seq (Figure 4E). n = 3 pairs, unpaired t-test, ns: not significant. **(G)** Latency in seconds for mice to move into the dark side of a shuttle box at baseline and 24 hours after training with shock to avoid that side. Data analyzed by one-way ANOVA with Sidak test for multiple comparisons, comparing HET to KO at baseline and at 24 hours; n=11 pairs, n.s.: not significant. **(H**,**I)** Number of lever presses on the last day (H) and time to learn task (I) by HET and KO mice in FR1 training to press a lever for food reward after food deprivation. n = 5 pairs (H, I). Log-rank (Mantel-Cox) test performed in (I) is not significant.

